# Hacking The Diversity Of SARS-CoV-2 And SARS-Like Coronaviruses In Human, Bat And Pangolin Populations

**DOI:** 10.1101/2020.11.24.391763

**Authors:** Nicholas J. Dimonaco, Mazdak Salavati, Barbara Shih

## Abstract

In 2019, a novel coronavirus, SARS-CoV-2/nCoV-19, emerged in Wuhan, China, and has been responsible for the current COVID-19 pandemic. The evolutionary origins of the virus remain elusive and understanding its complex mutational signatures could guide vaccine design and development. As part of the international “CoronaHack” in April 2020 (https://www.coronahack.co.uk/), we employed a collection of contemporary methodologies to compare the genomic sequences of coronaviruses isolated from human (SARS-CoV-2;n=163), bat (bat-CoV;n=215) and pangolin (pangolin-CoV;n=7) available in public repositories. Following *de novo* gene annotation prediction, analyses of gene-gene similarity network, codon usage bias and variant discovery were undertaken. Strong host-associated divergences were noted in ORF3a, ORF6, ORF7a, ORF8 and S, and in codon usage bias profiles. Lastly, we have characterised several high impact variants (inframe insertion/deletion or stop gain) in bat-CoV and pangolin-CoV populations, some of which are found in the same amino acid position and may be highlighting loci of potential functional relevance.

## 1. Background

The continued and increasing occurrence of pandemics that threaten worldwide public health due to human activity is considered to be inevitable [1,2]. The COVID-19 (2019-current) pandemic caused by the emergence in Hubei, China, of what has now been identified as Severe Acute Respiratory Syndrome Coronavirus 2/ Novel Coronavirus 2019 (SARS-CoV-2/2019-nCoV) by The Coronaviridae Study Group [3], has brought a number of questions regarding its transmission, containment and treatment to the urgent attention of researchers and clinicians. The urgency of such questions has spurred a number of atypical approaches and collaborations between experts of different fields and as such, this study was carried out as part of a “CoronaHack” hackathon event in April 2020 where the authors gained access to genomes and related metadata available at the time (Dec 2019 - April 2020).

Viruses of the Coronaviridae family have long been studied and while there have been great advances in our understanding, each new emergence has brought about its own questions The sub-family Coronavirus consists of four genera, Alphacoronavirus (Alpha-CoV), Betacoronavirus (Beta-CoV), Gammacoronavirus and DeltaCoronavirus. Coronaviruses are a family of single-stranded, enveloped and extremely diverse RNA viruses which have come into contact with humans numerous times over the past few decades alone [4]. At around 30kb, they exhibit at least six Open Reading Frames (ORFs), ORF1a/b comprising of approximately 2/3 of the genome which encodes up to 16 non-structural replicase proteins through ribosomal frame-shifting, and four structural proteins: membrane (M), nucleocapsid (N), envelope (E) and spike (S) glycoprotein [5]. Coronaviruses have developed a number of different strategies to infiltrate their host-cells. In human-associated CoVs, it has been shown that different parts of the human Angiotensin Converting Enzyme 2 (hACE2) can be bound to by their respective S proteins. Pathogens such as SARS-CoV-1 (Severe Acute Respiratory Syndrome Coronavirus) and MERS-CoV (Middle East Respiratory Syndrome Coronavirus) have shown Coronaviruses to be capable of presumed efficient adaptation to their human host and exhibit high levels of pathogenicity [6,7]. Interestingly, SARS-CoV-1 and MERS, which along with SARS-CoV-2 are both Beta-CoVs, exhibit only 79.5% and 50% sequence similarity at the whole genome level to SARS-CoV-2, whereas SARS-CoV-2-like coronaviruses found in pangolins (pangolin-CoVs) and bat coronavirus (bat-CoV) SARSr-Ra-BatCoV-RaTG13 (RaTG13) are 91.02% and 96% respectively [8]. The relationship of SARS-CoV-2 to other SARS-like coronaviruses, the possible role of bats and pangolins as reservoir species and the role of recombination in its emergence, are of great interest [9]. Speculations around other intermediary hosts are also at play, which might have affected the ability for zoonotic transmission for SARS-CoV-2 to its human host [10]. Crucially, this evolutionary relationship between SARS-CoV-2 and its lineage may prove to be an important factor in the eventual management or containment of the virus. Moreover, the mutation events along the evolutionary timeline of SARS-CoV-2 are of importance in the discovery of possible adaption signatures within the viral population. At the time of the hackathon, there were two main suspected SARS-like reservoir host species; bat and pangolin (named bat-CoV and pangolin-CoV).

With this in mind, our study aimed to systematically compare a broad selection of contemporary available SARS-CoV-2, bat-CoV and pangolin-CoV at genome, gene, codon usage and variant levels, without preference for strains or sub-genera. This was comprised of 46 SARS-CoV-2 genomes isolated early in the pandemic from Wuhan, China (Late 2019-Early 2020), 117 SARS-CoV-2 genomes isolated in Germany, representing the later stage of global transmission, 215 bat-CoV genomes of Alpha-CoVs and Beta-CoVs and 7 pangolin-CoV genomes, of which 5 were annotated as Beta-CoVs. During the hackathon, it was recognised that potential biases can arise from directly comparing SARS-CoV-2 to a wide repertoire of coronaviruses of varying stages of genome annotation. Therefore, we performed a new comparative annotation of all genomes used in this study. To further validate mutational adaptations which may have facilitated the zoonotic transmission of SARS-CoV-2, a codon usage analyse was carried out between the SARS-CoV-2 reference genes and the genes identified using the abovementioned approaches.

In addition, we profiled codon usage bias across our dataset, as in the process of host adaptation, viruses can evolve to express different preferential codon usages [11–13]

Through examining the inherent sequence diversity between a comprehensive collection of SARS-CoV-2, bat-CoV and pangolin-CoV, we aimed to highlight naturally occurring high impact variations that can potentially introduce a moderate change in the resulting protein, such as the insertion or deletion of an amino acid or early termination of the sequence. Understanding the stability and variability of these positions may potentially aid future design of vaccines or treatments. For instance, an amino acid position where insertion or deletion is commonly found in a coronavirus affecting other species may indicate that its alteration does not have a dramatic impact on the overall protein folding, or that the position is important for transmission to a new host.

Our work is differentiated by the way a systematic approach was used to process a non-selective group of these viral genomes from public repositories, prior to applying a wide range of contemporary methodologies and genomic knowledge that highlight the variations that exist between different host species.

## 2. Results

### 2.1. Data Collection and Phylogenetic Analysis

We were able to collate 215 bat-CoV genomes of varying families (Alphacoronaviruses and Betacoronaviruses) with only one exhibiting a small proportion or genomic uncertainty (presence of 0.45% ‘N’ nucleotide). However, only 7 pangolin-CoV genomes, of which 5 were annotated as Betacoronaviurs, were available at the start of this study. 3 pangolin-CoV genomes also contained levels of the ambiguous ‘N’ nucleotide, two of them at high levels (6.88 and 8.19%). A population of post-outbreak SARS-CoV-2 genomes from Charite [14], Germany, were also collated for further analysis. For the phylogenetic analysis, we examined the complete set of 269 genomes (7 pangolin, 47 Wuhan SARS-CoV-2 isolates (including 1 Ensembl Wuhan reference) and 215 bat). The phylogenetic tree produced at the whole genome level showed a clear separation between SARS-CoV-2 Wuhan isolates and the bat-CoV genomes (except RaTG13) (Figure 1). The 7 pangolin-CoV genomes cluster together and are closest to the SARS-CoV-2 Wuhan isolate population of genomes, discounting the RaTG13 genome which was the closest to SARS-CoV-2. The Ensembl Wuhan reference genome [15] has been placed within the other Wuhan isolates.

**Figure 1.**
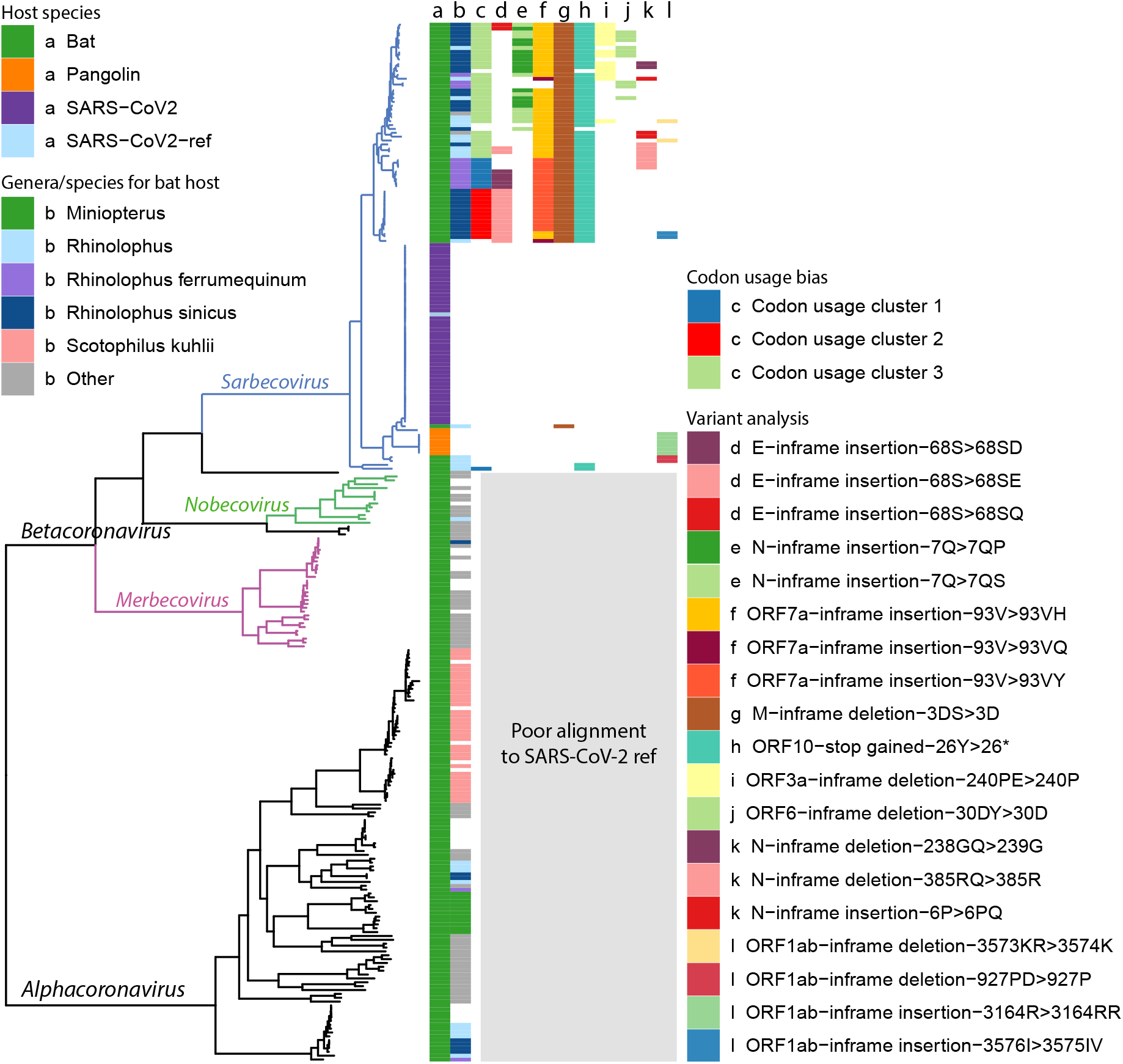
Ladderised phylogenetic tree of bat-CoV, pangolin-CoV and SARS-CoV-2 (Wuhan dataset and reference) genomes. Metadata are indicated on the top left corner, including a) dataset name and b) the bat genera and species if the genome is of bat host. Clades for Betacoronavirus subgenera, Sarbecovirus, Nobecovirus and Merbecovirus, are indicated on the graph, showing that our codon usage bias and variant analysis results are restricted to the Sarbecovirus due to poor alignment between SARS-CoV-2 ref and genomes outside this subgenera. There also appears to be some degree of genera and species separation for bat hosts. The majority of the Sarbecovirus affect the bat genus Rhinolophus (column b, light blue, dark blue and purple), whereas a much smaller proportion of the Alphacoronavirus are found in bats of this genus. Some clades overlap with specific bat species, including Rhinolophus ferrumequinum, Rhinolophus sinicus and Scotophilus kuhlii. The results from the analysis made in later parts of this study are also highlighted, including c) codon usage bias clusters, d-f) high impact variants with multiple variants are found in the same amino acid position, g-j) other high impact variants with a single amino acid change found in > 10 genomes, k-l) other high impact variants.

The tree produced was used as an analytical anchor for which we could use to refer to in the codon usage bias and variant analysis. High impact variants and codon usage clusters were plotted on the tree to show their distribution across the different clades along the topology of the tree.

### 2.2. Gene Identification

The complimentary approach of using PROKKA [16] and BLAST [17] to identify the set of genes for each of the viral genomes complemented each other and enabled a comparative analysis. A breakdown of the number of genes identified for each dataset is shown in Table 1 and Table A1 presented the number of genes annotated by PROKKA or BLAST.

**Table 1.** This table presents the distribution of the number of predicted genes for each dataset. Bat-CoV exhibit the widest distribution of gene count, and pangolin-CoV has the highest number of gene count, with one genome having 17 predicted genes. These outliers have low sequence or assembly quality. In the case of the pangolin-CoV genome reporting 17 genes, it has low quality (‘NNNN’) nucleotide regions spanning the centre of genes, which causes PROKKA to identify the two ends of one gene. The median gene count only varying in bat-CoVs, likely attributed to the large phylogenetic variation exhibited across the bat genomes.

| Dataset | Min. | Median | Mean | Max. | Sample Count |
| --- | --- | --- | --- | --- | --- |
| Wuhan | 7 | 11 | 11 | 13 | 46 |
| Charite | 9 | 11 | 11 | 12 | 117 |
| Bat | 2 | 9 | 9 | 12 | 215 |
| Pangolin | 10 | 11 | 12 | 17 | 7 |

BLAST was utilised in attempts to capture a number of genes with strong homology to the SARS-CoV-2 ref (≥ 80%) that were not identified by PROKKA. In particular, these genes were E, ORF8 and ORF10.

Whilst this has enabled the characterisation of E and ORF10 in many genomes, no additional ORF8 were identified through BLAST apart from 6 examples in the Charite dataset (genomes had levels of ambiguous ‘N’). This could in part be due to the high threshold setting used in the BLAST search (>80% identity). ORF8 was only identified in 3 bat-CoVs and 1 pangolin-CoV with this combined approach. At least 38 additional bat-CoV ORF8 and 4 additional pangolin-CoV ORF8 representatives were identified by PROKKA with less than 80% identity. Genes utilising ribosomal frameshifting such as the aforementioned ORF1ab, are inherently difficult to identify correctly without extensive analysis involving techniques and evidence such as RNA expression analysis. For the majority of genomes studied, PROKKA was able to identify two large ORFs spanning almost the entire length of the ORF1ab locus and detect a central coronavirus frame-shifting stimulation element (named Corona_FSE and separating the two ORFs) which is a conserved stem-loop of RNA found in coronaviruses that can promote ribosomal frameshifting [18]. The gene sequences generated by PROKKA and BLAST (E and ORF10) were used for downstream analysis, including gene-gene network graph, codon usage bias analysis, and a gene-presence summary table. The gene-presence summary table notates whether SARS-CoV ref genes were found (≥ 80% and ≥ 50% sequence coverage) in each genome; this table is available in the GitHub project https://github.com/coronahack2020/final_paper/tree/master/host-data. Supplementary files for each host (in each folder) are named as *\*_genome_metrics.csv.*

### 2.3. Gene Relationship Network Graph

A gene-gene similarity network analysis was used to compare genes across SARS-CoV-2, bat-CoV and pangolin-CoV. The advantage of using a 3D network approach to visualise this information was that it simplifies complex information as patterns. Genes sharing high similarity form independent clusters. In cases where there is a high degree of dissimilarity in a gene for different host species, a pattern of 2 or more distinct clusters would take place, with each cluster comprised of genes derived from samples of the same host-species. In genes where there is a medium level of dissimilarity across host-species, two or more cluster would appear fused and potentially break apart into distinct clusters if the edge threshold were increased. Both of these patterns are observed within this dataset. Distinct separation by host species are seen in ORF1a, ORF3a,ORF6, ORF7a, ORF8 and S (Figure 2). The strongest host-species separation observed were between SARS-CoV-2 and bat-CoV; pangolin-CoV always group closer to SARS-CoV-2 than to bat-CoV. In the cases of ORF3a, ORF8 and S, complete separation was observed between bat-CoV and human SARS-CoV-2 (Figure 2B & C). One bat-CoV genome, RaTG13, was more similar to SARS-CoV-2 and pangolin-CoV than the remainder of the bat-CoV for S (Figure 2C). For ORF3a, three bat genomes (MG772933, MG772934 and MN996532; named bat-SL-CoVZC45, bat-SL-CoVZXC21 and RaTG13 respectively) clustered together with SARS-CoV-2 and pangolin-CoV rather than with the remainder of the bat genomes (Figure 2). These same three genomes are the only bat-CoV with ORF8 that co-cluster with SARS-CoV-2 ORF8 under the percentage identity threshold (≥80%) set for building the network graph. Other bat-CoV ORF8 were so distinct from SARS-CoV-2 ORF8 that they do not co-cluster, even when edge filtering was removed.

**Figure 2.**
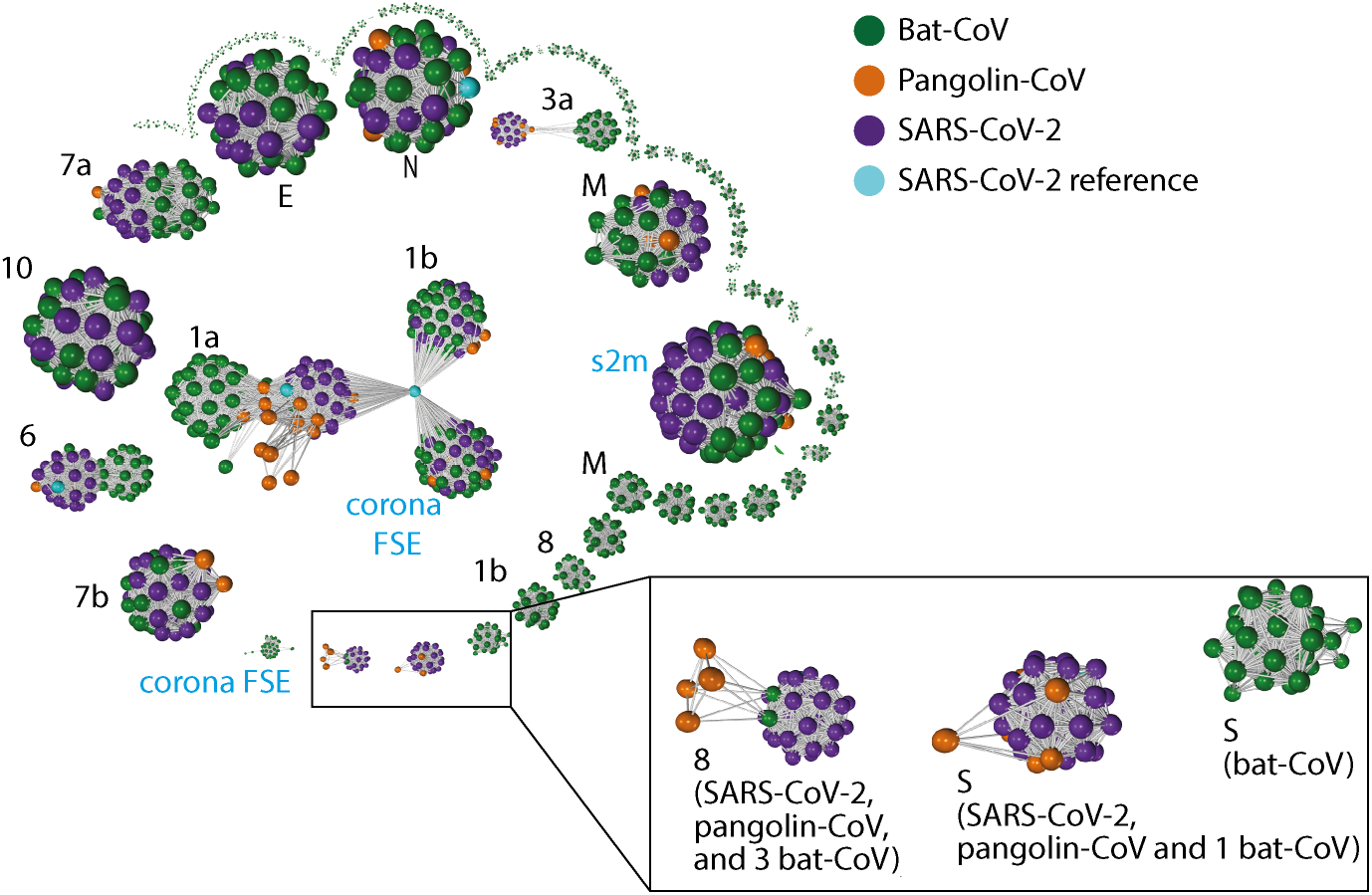
Gene-gene similarity network analysis. Each node represents a gene defined by PROKKA or a DNA segment similar to genes from the SARS-CoV-2 reference genome. The nodes were compared against each other using BLAST, and nodes with high similarity (BLAST score ≥ 60 and a query coverage ≥ 80%) were connected with an edge. The network graph is labelled with host species. The black font in the graph indicates the corresponding SARS-CoV-2 gene names (“ORF” omitted) for the larger clusters, whereas blue font indicate additional non-coding sequences defined by PROKKA. Instead of the full length ORF1ab (21k in length), ORF1a and ORF1b were defined by PROKKA as two separate genes. Notably ORF1a, ORF3a, ORF6, and ORF8 and S, show strong separations between nodes from different species. ORF8 from 3 bat-CoV co-cluster with ORF8 from SARS-CoV-2 (RaTG13, bat-SL-CoVZC45 and bat-SL-CoVZXC21 respectively). The remaining bat-CoV ORF8 do not co-cluster with SARS-CoV-2 ORF8 even without the edge filtering threshold. For S, the bat-CoV RaTG13 co-cluster with COVID-19 and pangolin. A cluster of bat-CoVs break off for ORF1b and M, suggesting a large amount of variation amongst bat-CoV for these genes.

To investigate whether if potential gene transfer or recombination that may have come from more distantly related bat-CoV, we sought for unusual co-clustering between genes characterised from bat-CoV and SARS-CoV-2. We did not observe such pattern; RatG13 co-cluster with SARS-CoV-2 for many genes, but it is also the most similar bat-CoV to SARS-CoV-2 at a genome level.

Two additional genes identified by PROKKA, Corona FSE, a non-coding frame-shift stimulation element within ORF1ab and s2m, a stem-loop II-like motif [19] have both been shown to be highly conserved and important for SARS-2-like coronaviruses. s2m has been identified as a mobile genetic element which has been described in a number of single-stranded RNA virus and insect families and has also been shown to be important for viral function [20,21].

In summary, the use of gene-gene network analysis enables us to determine groups of closely related genes, which not only highlights genes showing strong host-species separation, but also characterise clusters of related genes that may be absent or highly different from the reference genome of interest, such as ORF8. 6 genes, ORF1ab, ORF3, ORF6, ORF7a, ORF8 and S, showed a strong host-species separation in the network graph. In particular, with the exception of S, where bat-SL-CoVZC45, bat-SL-CoVZXC21 clustered closer to bat-CoVs, the bat genomes, bat-SL-CoVZC45, bat-SL-CoVZXC21 and RaTG13, clustered together with SARS-CoV-2 than the remainder of the bat-CoV for these 5 genes.

### 2.4. RNAseq expression analysis

In our exploratory analysis during the Hackathon event, we attempted to capture gene-level expression evidence for each of the predicted ORFs. However, following the event, we recognise that RNA virus gene expression cannot be captured through standard RNAseq analysis pipeline. We have included the results of our analysis in the supplementary section for record purpose only; it is an inaccurate estimation of the viral gene expression, as it does not differentiate viral mRNA expression from viral genome.

### 2.5. Codon Usage Bias

Codon usage profiling of all representative genes of the SARS-CoV-2 ref separated from human host (Wuhan and Charite datasets), bat-CoV and pangolin-CoV was carried out. RSCU (relative synonymous codon usage) were calculated for each gene and for all genes that are found in >18% of the bat dataset s (E, N, S, ORF1a, ORF3a and ORF10) to depict an overall relative synonymous codon usage across genomes from the datasets. Principle component analysis (PCA) using RSCU showed a strong host-species separation; the first principle component (PC1) accounts for > 90% of the variation (Figure 3a and b). Some separation was observed amongst bat-CoVs (Figure 3a & b). K-means clustering was used to cluster bat-CoVs using the multiple-gene PCA output (with the exception of MG772933, MG772934 and MN996532, named bat-SL-CoVZC45, bat-SL-CoVZXC21 and RaTG13 respectively, as they group closer to SARS-CoV-2 and pangolin-CoV). The generated clusters, unsurprisingly correspond to different clades in the phylogenetic tree (Figure 3b and Figure 1c). We have also examined RSCU across bat-CoV, pangolin-CoV and SARS-CoV for each gene. Strong host-species separation is seen across all genes. Similar to the PCA done with multiple genes, whilst the majority of the variation can be explained by host-species differences, there is also some variation amongst the bat-Cov that correspond to the k-means clusters generated from the multi-gene PCA analysis (Figure A3). A summary of the synonymous codon ratios (the number of codon divided by total number of codons coding for the same amino acid), sorted by amino acids, are shown in Figure A5.

### 2.6. Variant Analysis

Haplotype aware variant calling and variant effect prediction of all genomes in the study has been summarised in Figure 4 and Table additional file 1. There are a total of 1,127 variants that are missense, inframe deletion, inframe insertion, stop gained, stop lost, as can be seen in Figure A6. We have removed missense from further analysis and came to a total of 24 high impact variations in 8 genes were when comparing bat-CoV and pangolin-CoV genomes against the SARS-CoV-2 ref. We have annotated the majority (with the exception of the NC045512_27675A>ACAG) of these variation in Figure 1, and found that some of these variations, such as variants identified in E, ORF7a and ORF3a, appear to exhibit some degree of clade specificity. The only stop gain variant (i.e. NC045512_29635) was present in ORF10 gene of 57 bat-CoV genomes (29635 bp position C>A) which was only representing a synonymous variant in the same position of 6 pangolin-CoV genomes. This variant affected 26Y>26* (Tyrosine to STOP codon TAC>TAA) in bat ORF10. Assuming the direction of host-selection from bat and pangolin to human, this variant could explain the presence of a longer ORF10 isoform in the 2 latter hosts in comparison to bat-CoV. From the variant Table 4, four in-frame insertions were identified as follows:

- ORF1ab gene at position 9757 (NC045512_9757 T>TAGA 3164R>3164RR) of all pangolin-Cov genomes which represents an extra Arginine.
- E gene at position 26448 (NC045512_26448 T>TGAA 68S>68SE) in 33 bat-Cov genomes which caused an addition of Glutamine.
- ORF7a gene at position 27672 (NC045512_27672 T>TCAC 93V>93VH) in 24 bat-Cov genomes by addition of an Histamine.
- N gene at position 28293 (NC405512z_28293 A>AACC 7Q>7QP) in 13 bat-Cov genomes by addition of a Proline.

**Figure 3.**
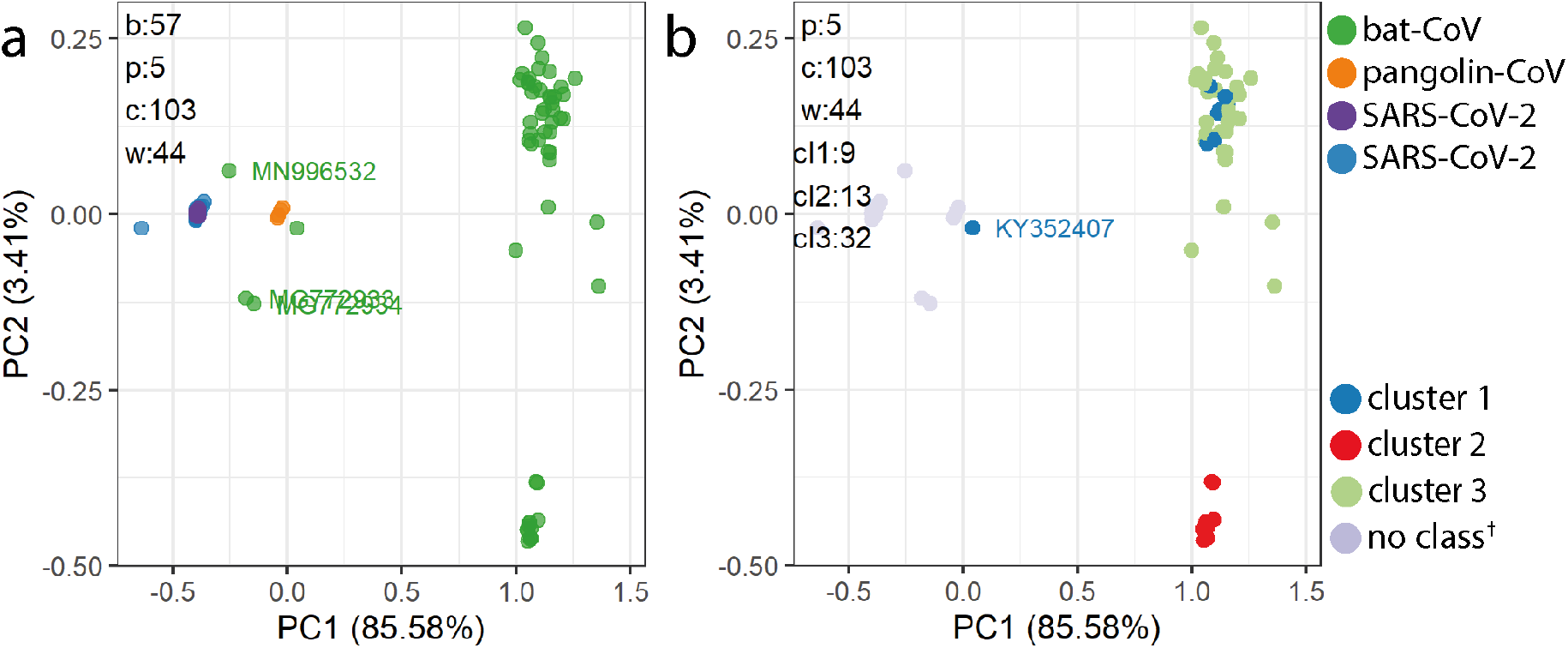
A) Principle component analysis was carried out on the relative synonymous codon usage (RSCU) to demonstrate host-species variation in codon usage bias. RSCU was calculated using E, N, S, ORF1ab, ORF3a and ORF10. This analysis only included genomes annotated with all of the listed genes. The majority of the variation (principle component, PC1) can be explained by host-species, with the exception of the three labelled bat-CoV (MG772933, MG772934 and MN996532; named bat-SL-CoVZC45, bat-SL-CoVZXC21 and RaTG13 respectively). B) By using k-means clustering, two distinct clusters were generated from bat-CoV (excluding the 3 distinct bat-CoV highlight in a). A portion of the variation (PC2) reflect the different clades in the phylogenetic tree (Figure 1). Both cluster 1 and cluster 2 remains grouped closely together across all genes (with the exception of KY352407. Figure A3).

**Figure 4.**
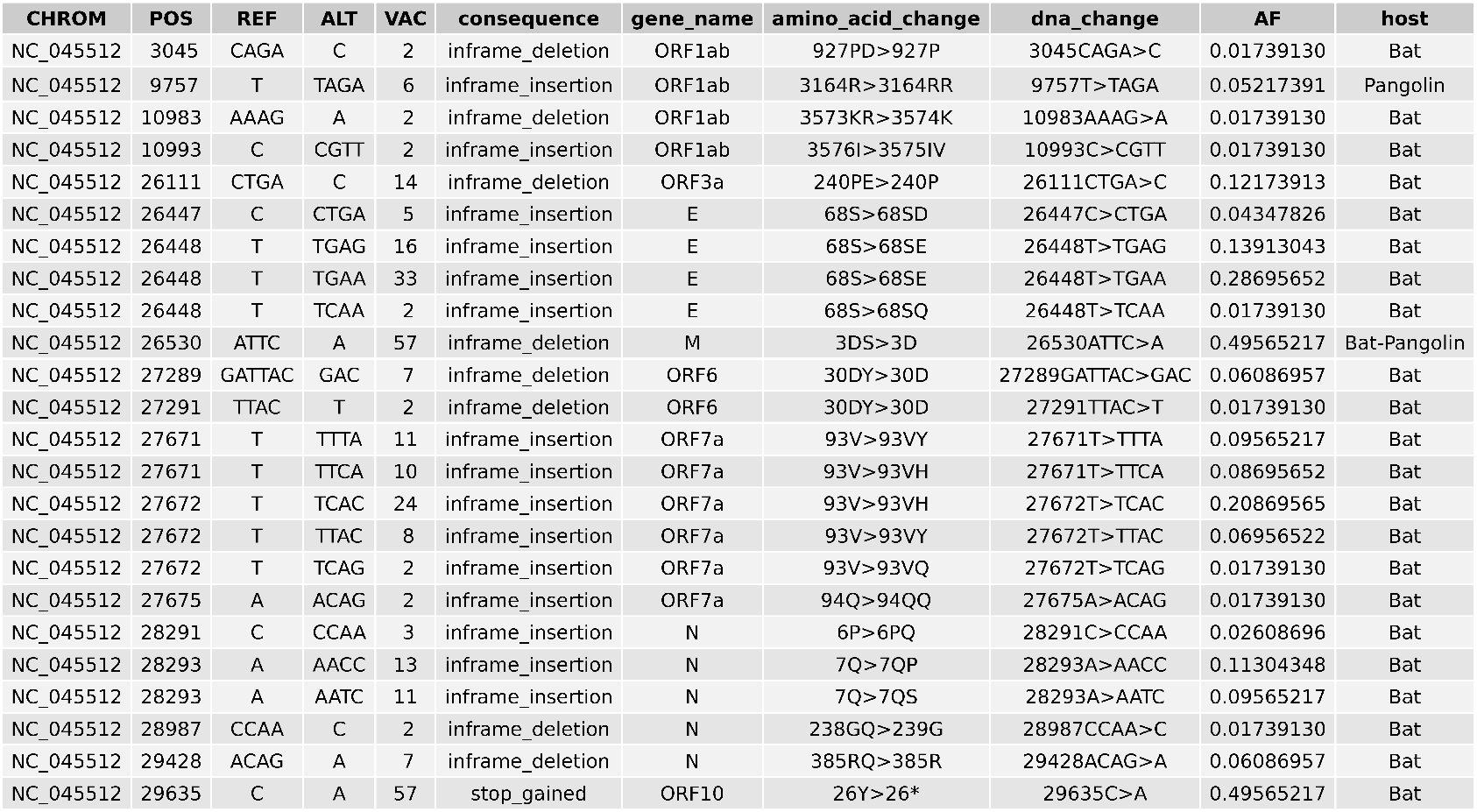
High impact variants identifies across bat and pangolin genomes using the variant calling pipeline based on SARS-Cov-2 Ensembl reference genome.

Two in-frame deletions were also identified in ORF3a and M genes. A single Glutamine deletion in ORF3a at position 26,111 was present in 14 bat-Cov genomes (NC045512_26111 CTGA>C 240PE>240P) and a Serine deletion in M gene at position 26,530 (NC045512_26530 ATTC>A 3DS>3D) was present in 57 bat-Cov genomes.The same position showed a missense mutation of 3D>3A (in 2 bat-Cov [bat-SL-CoVZC45 and bat-SL-CoVZXC21] and 1 pangolin-Cov) and 3D>3G in 6 pangolin-Cov genomes.

## 3. Discussion

During the 5 day hackathon, we endeavoured to utilise the genomic data aggregated by the scientific community and undertook a multifaceted and comprehensive exploration of the genomic sequences (or “similarities and differences”) of coronaviruses infecting bat and pangolin hosts, available at the time. We have compared SARS-Cov-2 to all bat-CoV and pangolin-CoV genomes from the listed data repositories (NCBI, VIPR and Databiology) without selecting for strains to represent any specific genera, species or sub-strain. Our comparisons spanned across several levels: whole-genome, genes, codons and individual variants.

The phylogenetic tree inferred from all genomes studied in this manuscript presents a picture of vast bat-CoV diversity and its topology is similar to those of previous studies carried out on pangolin and bat coronaviruses when compared to the SARS-CoV-2 genome [22]. Previous phylogenetic profiling has noted that RaTG13 (bat-CoV) bares the closest resemblance to SARS-CoV-2 using 14 SARS-CoV-2 and 55 non-SARS-CoV-2 coronavirus genomes [23]. In this study, we have investigated a more expansive set of bat-CoV genomes, and included pangolin-CoV genomes. RaTG13 remains the closest to SARS-CoV-2 at the whole-genome level, although all 7 pangolin-CoV genomes are more closely related to SARS-CoV-2 than the remaining 214 bat-coronavirus (Figure 1). This relationship has previously been reported and a recombination event between pangolin-CoVs and RaTG13 has been theorised [24]. The RaTG13 coronavirus found in horseshoe bats, as with SARS-CoV-2, is a member of the coronaviridae subgenus Sarbecovirus, has been suggested to be the closest relative to SARS-2 in a number of studies [25]. The origin of SARS-CoV-2 is still unknown and a number of coronaviruses from different hosts have been proposed [26,27]. Bats are often linked to SARS-like viruses capable of zoonotic host transfer due to their unique niche as viral reservoirs, meaning that they are relatively unaffected by viral loads and their natural proximity to human habitation [28,29]. Recombination has been suggested as an avenue for host-transfer for a number of RNA viruses such as SARS-CoV-1 and MERS [30]. More recently, evidence has also been found for inter-host recombination events in a SARS-CoV-2 patient, which may have lead to new traits such as increased virulence from multiple strains [31].

In the attempts to address the potential recombination events or gene transfers between a strain distantly related to SARS-CoV-2 and a strain more closely related to SARS-CoV-2, we sought to annotate, characterise and compare genes from our diverse sets of coronaviruses. RNA virus genomes are often compact, with little intergenic distance between genes, even those of the Coronaviridae family which are regarded to have the largest RNA viral genomes. This makes accurate annotation a difficult task, especially for frame-shift utilising genes and the distinction of what is produced as final protein product. We initially encountered a number of problems while performing genome annotation, where a number of contemporary gene prediction methodologies failed to identify ORF10 in any of our datasets, except for 5 pangolin-CoV genomes. Furthermore, the DNA sequence representing ORF10 in SARS-CoV-2, which was previously reported as having no homology to any known sequence in public databases [32], has now been found in previous examples of coronaviruses infecting both pangolins and bats with very high sequence similarity (≥90%) [33]. With the utilisation of BLAST, ORF10 was found in 162 out of the 163 of all the SARS-CoV-2 (1 genome contained low quality regions), all pangolin-CoVs and 59 bat-CoV genomes. On the other hand, we initially found only 3 bat-CoV (RaTG13, bat-SL-CoVZC45 and bat-SL-CoVZXC21) ORF8 representatives when comparing PROKKA characterised sequences against SARS-CoV-2 genes and identified no additional sequences through BLAST. However, with the use of gene-gene network analysis, we noted that this apparent absence of ORF8 was due to the very low percentage similarity between most bat-CoV ORF8 and SARS-CoV-2 ORF8; the network analysis showed a cluster of 38 bat-CoV ORF8 that strongly correlated to each other. Ceraolo et al. (2020) have shown that ORF8 from RaTG13 shares 94% protein identity to SARS-CoV-2, whilst those of other bat-betacoronaviruses show <60% similarity [34]. Furthermore, Pereira (2020) has shown ORF8 orthologues are present outside betacoronaviruses linage B (subgenus Sarbecovirus) [35]. Interestingly, the 3 bat-CoV ORF8 genes were more similar to SARS-CoV-2 than the majority of the pangolin-CoV ORF8 representatives; 4 of the 5 pangolin-CoV ORF8 genes only joined to the ORF8 cluster through the bat-CoV ORF8 in our network analysis. There were only 4 proteins with 80-100% identity and 100% coverage identified by BLAST searches against ORF8 using the NCBI and UniProtKB/Protein databases [36]. Two of these proteins were bat-CoV (RaTG13 and Bat-SL-CoVZC45), while the other two were pangolin-CoV; all four genomes were present in our dataset and we have also observed this high similarity in ORF8. The exact function of ORF8 remains to be elucidated, although studies on ORF8 from SARS-CoV-2 and ORF8ab and ORF8b from SARS-CoV-1 have suggested a role in immune modulation through the interferon signalling pathway [37,38] and induce strong antigen response [39]. Although the origin or function of the SARS-related coronavirus ORF8 remains unresolved, a 29-nucleotide deletion in ORF8 is often found in SARS-CoV-1, when compared to civet-CoV, suggesting that ORF8 may be important for interspecies transmission [40]. In post-pandemic studies of the SARS-CoV-1 coronavirus, deletions in specific genome domains found in samples from human and mammalian hosts were identified as being possible conduits for early human infection [41].

Other genes that show strong host-species separation in the gene-gene network analysis include ORF1a, ORF3a, ORF6 and S. In contrast to ORF8, where the three bat-CoV were more similar to SARS-CoV-2 than pangolin-CoV, pangolin-CoV and SARS-CoV-2 S protein were more similar to each other (97.5%), than those of RaTG13 and SARS-CoV-2 (95.4%) [33]. This is significant as the S protein plays an important role in the initial penetration and infection of host cells [42]. Several human coronaviruses, including SARS-CoV-2, SARS-CoV-1 and human coronavirus NL63 (hCoV-NL63), enters the host cells by binding to the host cell angiotensin-converting enzyme 2 (ACE2) through the receptor binding domain (RBD) of S protein [43,44]. Host-cell receptor recognition is one of the determining factors of host-cell tropism and the co-evolutionary struggle between viruses and their hosts has likely involved a number of exchanges of genetic information during long periods of interaction of pathogen and host-cell contact [45,46]. Viruses have been shown to have high degrees of flexibility in their receptor usage and poses capacity to reach efficient binding through mutations [45,46]. By altering the amino acids within the RBD of SARS-CoV-1, Qu et al. (2005) has noted that a single amino acid substitution reduces the binding affinity, and two amino acid substitution almost abolishes its infection of human cells [47]. Moreover, by substituting these amino acids civet-CoV for those from SARS-CoV-1 enabled the modified civet-CoV to infect human ACE-2 expressing cells [47]. This illustrates the importance and complexity of S in cross-species infectivity. Nonetheless, it would appear that despite the S protein being more similar between pangolin-CoVs and SARS-CoV-2, as compared to SARS-CoV-2 versus bat-CoVs, the S protein in RaTG13 on the whole is still more similar to that of SARS-CoV-2 than to those of all other bat-CoVs in this study (Figure 2C). This supports the theory that neither a currently sequenced pangolin-CoV or bat-CoV are the most recent ancestor of SARS-CoV-2.

In addition to examining the overall sequence similarity of between genes derived from bat-CoV, pangolin-CoV and SARS-CoV-2, we have also examined the codon usage within and across genes. Codon usage bias across the species-host range may show signs of preferential codon mutation which have occurred during the complex process of host interaction and transfer [11,12]. The knowledge of nucleotide profiles and subsequent codons during the human-virus co-evolution could be invaluable to the design of vaccines and their continuous development over the years to come [48]. We have demonstrated a strong host-species separation in the overall codon usage when combining multiple genes (E, N, S, ORF1a, ORF3a and ORF10) in the analysis. There is very little variation in codon usage bias within the SARS-CoV-2 isolates. However, all pangolin-CoVs and the 3 bat-CoVs (bat-SL-CoVZC45 and bat-SL-CoVZXC21 and RatG13) have a more similar codon usage to SARS-CoV-2. The k-means clusters generated from the PCA using RSCU of multiple genes correspond to clades within the phylogenetic trees and remain intact when compared across each gene individually (Figure 1c and Figure A3), with two clusters aligned with subsets of bat-CoVs isolated from Rhinolophus ferrumequinum and Rhinolophus sinicus respectively. When comparing codon usage bias across the host-species at a gene-level, bat-CoV also appear to be more distinct from SARS-CoV-2 than pangolin-CoV, both with respect to the percentage similarity and the presence/absence of genes, with the exception of the 3 bat-CoVs (bat-SL-CoVZC45, bat-SL-CoVZXC21, and RaTG13). In contrast to the codon usage analysis carried out by Gu et al. (2020), in which the authors has reported the codon usage for M in pangolin-CoVs to be more similar to those of SARS-CoV-2 than RatG13 [49], our analysis does not suggest this to be the case. This could be due to a difference in the range of hosts included; we have included SARS-CoV-2, pangolin-CoV and bat-CoV, whereas they additionally included coronavirus that affects camel, rodent, pigs and other species. Our codon usage analysis has been restricted to an overall comparison of RSCU across the genomes we have used in this study, as more detailed breakdown of codon usage bias and CpG dinucleotide have been carried out elsewhere [50–52]. Previous studies have correlated the RSCU of SARS-CoV-2 to those of human genes and found them to significantly correlate with a large number of human genes, which are enriched in pathways relating to host response to viral infection [50]. It has been observed that host genes sharing similar codon usage as SARS-CoV-2 are downregulated during an infection, potentially through causing an unbalance to the host tRNA pool and thus host protein synthesis [51]. These mechanisms potentially reflect the affects of coronavirus on the genome separation of different host species, observed in the RSCU analysis.

Next, we focused on variants that could potentially have a more profound impact on the structures of the proteins through the addition or removal of an amino acid, or through early termination. In this analysis, we have found that only pangolin-CoV and a subset of bat-CoV (Sarbecovirus or unannotated) were similar enough to the SARS-CoV-2 ref for the sequences to align 1. Population level viral mutation is a complex process, involving a number of pressures, and while RNA viruses often exhibit some of the highest mutation rates of all viruses, conserved variants can exhibit important functional changes such as the ability to evade immunity more efficiently [53]. Unlike the vast majority of RNA viruses, coronaviruses encode a complex RNA-dependent RNA polymerase that has a 3’ exonuclease domain [54], effectively proofreading mutational events and therefore are less error-prone. Therefore the mutations observed across populations have undergone an error-correction process which means they are more likely to be functionally beneficial to the virus. We have observed several of such variants that are at consistent loci across different bat-CoV clades as shown in Figure 1. Some of these variants are seen in the majority of the bat-CoV samples (which align to SARS-CoV-2 ref), including a stop-gain for ORF10 and an inframe deletion for M, whilst others, such as the variants seen in ORF7a and E appear to be clade specific (Figure 1). Several of these variants affect the same amino acid positions, including E (inframe insertion of *Asp* (Aspartic acid), *Glu* (Glutamic acid) or *Gln* (Glutamine) at at positions 68), N (inframe insertion of *Pro* (Proline) or *Ser* (Serine) at position 7) and ORF7a (inframe insertion of *His* Histidine, *Gln* or *Tyr* (Tyrosine) at position 93) (Figure 1). Notably, the stop-gain was identified at amino acid position 26 in ORF10 for 57 of the 59 bat-CoV genomes with ORF10 that had >80% similarity to the SARS-CoV-2 ref. The absence of this stop codon in the pangolin (which exhibited synonymous mutations at the same locus) and human adapted viruses could result in a longer isoform of the ORF10 or fundamental changes in its function and expression levels. In a previous study of SARS-CoV-2 and pangolin-CoV genomes, position 26 was also identified as a region of population level variation from *Tyr* and *His* which significantly modifies the secondary structure of the coil region of the protein [55].

There has been little research on ORF10 function, and its expression has been the subject of debate. Whilst Kim et al. (2020) found little evidence of ORF10 expression (0.000009% of viral junction-spanning reads) in cell culture (Vero cells) [56], Liu et al (2020) found it to be abundantly expressed in severe COVID-19 patient cases but barely detectable in moderate cases [57]. Discrepancies in ORF10 expression may be due to differences in the level of infection and host cell-type used in the studies, however the variants noted show potential functions due to host-species-level conservation.

Multiple codon insertions and deletions also exist in ORF1ab of pangolin-CoV and bat-CoV genomes, which with the polypeptide coding potential of the gene which covers 2/3 of the genome, is likely to impact a number of important and complex elements of the virus. Machinery needed for viral replication and the proofreading subunit required to safeguard coronavirus replication fidelity, are just two functions of the 16 polypeptides which form after the processing of ORF1ab, and therefore potentially include several key targets for antiviral drug development [58].

As opposed to the single ORF10 variant that is observed in the majority of the bat-CoV, we have observed 3 different amino acid insertions (4 different nucleotide changes) at position 68 of E in 4 different clades of bat-CoVs. The small envelope E protein is the smallest of coronaviruses’ major structural proteins, but also one of the least described [59]. E has been shown to be highly expressed inside infected cells and the viruses which are formed without E exhibit reduced levels of viral maturation and tropism. Expression of the E product was essential for virus release and spread, thus demonstrating the importance of E in virus infection and therefore vaccine development [60]. The 68th amino acid position we highlight in this study is in the c-terminal domain, which coincides with the previously reported motif in SARS-CoV-1 (also at 68th amino acid position) that binds to the host cell PALS1 protein to facilitate infection [61].

Less than 0.5% of 3,617 SARS-CoV-2 genomes have been found to have non-synonymous mutation in E, and of these, 20% are at the 68th amino acid position [62]. These changes in amino acid may alter the hydrophobicity at the locus, thus possibly influencing the protein functions and interactions [62]. Two of the E variants we highlighted use different codons for the same amino acid (GAG or GAA for *Glu*), which potentially suggests interplay between the selection pressures of codon optimisation and amino acid insertion into the protein product.

We have characterised a number of inframe insertions at the amino acid position 93 in ORF7a across 55 bat-CoV genomes, and at position 94 reported in 2. As with position 68 in E, position 93 in ORF7a has multiple codon insertions coding for the same amino acid but in two groups. In these two groups of bat-CoVs, an additional *His* is encoded for by two different codons and secondly, so is *Tyr* in another group. Intriguingly, ORF7a in SARS-CoV-1 has been shown to regulate the bone marrow stromal antigen 2 which inhibits the release of virions of human infecting viruses [63].

N is another gene for which we have shown multiple inframe insertion variants for the same amino acid position. The N protein is highly expressed during an infection, and plays a key role in promoting viral RNA synthesis and incorporating genomic RNA into progeny viral particles [64]. In gene N, we observed two inframe insertions at amino acid position 7 for *Ser* or *Pro* from two groups of bat-CoVs (13 and 11 respectively), as well as two inframe deletions at positions 238 and 385. For M in 57 bat-CoV and pangolin-CoV, there is an inframe deletion at position 3, which removed the amino acid *Ser*. At this amino acid position, a missense mutation of (Asp) to *Arg* is seen in 2 bat-CoV (bat-SL-CoVZC45 and bat-SL-CoVZXC21) and 1 pangolin-Cov, and (Asp) to Glycine (Gly) in 6 pangolin-Cov genomes. These same two bat-CoV have been shown to be more similar to SARS-CoV-2 than other bat-CoV on other comparative metrics. M plays an important role in its interactions with both E and S to incorporate virions into the host-cells, thus any mutation in either gene may cause a number of important changes across all.

As opposed to the majority of the identified variants, ORF6 only exhibits 2 different inframe deletions in position 30, which remove the same amino acid *Tyr*.

The amino acid positions we have highlighted through our variant analysis may constitute important differences in the function or folding potential of the protein product. We have summarised these in Figure 1. The naturally occurring variants we observed across bat-CoV and pangolin-CoV may be associated with selection advantage, such as virulence or the efficacy to infect a specific host species.

Weber et al. (2020) have interrogated 572 SARS-CoV-2 genomes isolated worldwide and characterised 10 distinct mutation hotspots that have been found in up to 80% of the viral genomes they examined [65]. Whilst our reported amino acid positions do not coincide with the 10 hospots they have reported, some of the genomes they examined display changes on or adjacent to our reported positions

Through employing a number of genomic analysis methodologies, this study has aimed to bring understanding of the diversity across SARS-CoV-2 and SARS-CoV-2-like coronaviruses by comparing a wide selection of available genomes from the starting point of the pandemic. We have highlighted a high degree of host-specices separation in ORF3a, ORF6, ORF7a, ORF8 and S, as well as in codon usage. A number of amino acid positions that demonstrate high impact variants (inframe insertion/deletion or stop gain) have also been identified in various bat-CoV and pangolin-CoV; these are potentially functionally important positions of the protein and warrants further research.

## 4. Methods

### 4.1. Genomes

Historically, genomes held in public databases have been fragmentary, resulting in multiple collections with overlapping examples with alternative naming schemes and annotations. Fortunately, a large collection of virus genomes of the Coronaviridae family (Coronavirus) deposited in databases such as the Virus Pathogen Resource (ViPR) [66] have been provided with both genomic sequence and metadata which has been examined for redundancy and comparative annotation. Coronavirus genomes isolated from humans, bats and pangolins used in this study were collected from multiple repositories and grouped by their host and source. The databases and groups are listed in Table 2.

**Table 2.**
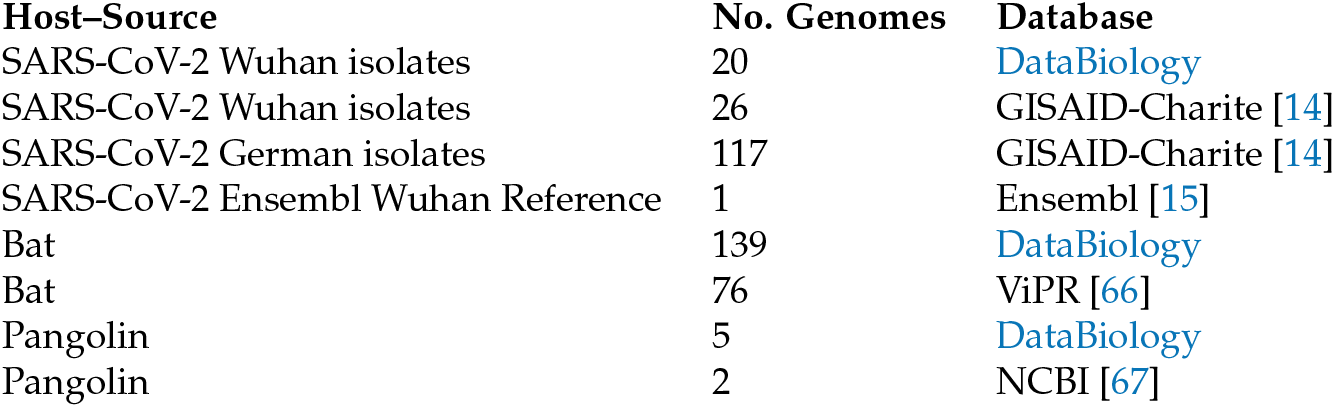
Coronavirus genomes were collected from the various database resources listed by host and source categories. Using taxonomic data made available by the Virus Pathogen Database and Analysis Resource (ViPR) [66], 70 bat-CoVs were identified as *Betacoronavirus* and 84 were *Alphacoronavirus.* 5 pangolin-CoVs were identified as *Betacoronavirus.* The remaining bat-CoV and pangolin-CoV genomes did not have a family identification. These were downloaded in May 2020 and consisted of the contemporary available and open datasets at the time. All genomes and their respective ID’s are currently available through NCBI (Oct 2020). In cases where two groups contained the same genome (Possibly with a different name), only one representative was taken.

| Host-Source | No. Genomes | Database |
| --- | --- | --- |
| SARS-CoV-2 Wuhan isolates | 20 | DataBiology |
| SARS-CoV-2 Wuhan isolates | 26 | GISAID-Charite [14] |
| SARS-CoV-2 German isolates | 117 | GISAID-Charite [14] |
| SARS-CoV-2 Ensembl Wuhan Reference | 1 | Ensembl [15] |
| Bat | 139 | DataBiology |
| Bat | 76 | ViPR [66] |
| Pangolin | 5 | DataBiology |
| Pangolin | 2 | NCBI [67] |

### 4.2. Genome Annotation

RNA viruses such as SARS and other coronaviruses have been characterised as having the ability to utilise ribosomal programmed frameshifting for a number of important genes [68]. Identification of such genes is complex and often requires high quality RNA expression evidence. Due to this and the complexity of genome annotation, especially in novel viral genomes such as SARS-CoV-2, two approaches were taken to identify the set of genes for each of the genomes in this study. In this regard, for defining genes, we first employed PROKKA (Rapid Prokaryotic Genome Annotation) to curate the genes for each of the coronavirus genomes. PROKKA utilises Prodigal [69] to initially find ORFs, which ensures that the DNA sequences of the genes found are in-frame and contain the correct amino acid coding potential. Prodigal is an unsupervised *ab initio* prediction method and therefore does not rely on previous knowledge to predict ORFs, which, unlike sequence homology based tools such as BLAST, does not require previously annotated sequence data to identify potential genes within novel genomes. Howeber, to overcome the limitations and intricacies of contemporary *ab initio* genome annotation techniques, BLAST was used to identify additional genes with strong homology to those present in the SARS-CoV-2 reference genome released by Ensembl v100 (SARS-CoV-2 ref) *ASM985889v3* [15](https://covid-19.ensembl.org). The additional BLAST annotation was performed with a BLAST percentage identity threshold of ≥ 80% are labelled separately where annotation methodologies may have an impact. This combined approach was used to avoid solely relying on either method, especially BLAST’s agnostic approach to coding frame detection.

### 4.3. Phylogenetic Trees

A Phylogenetic tree was produced from the genomes of the SARS-CoV-2 Wuhan isolates, Ensembl Wuhan reference and the bat and pangolin coronaviruses to examine their evolutionary relationships at the genomic level. Clustal Omega 1.2.4 [70] was used to perform a multiple sequence alignment for each of the genomes with default parameters. The phylogenetic tree was inferred from the multiple sequence alignment with RAxML [71] using default parameters apart from the GTRGAMMA option and bootstrapping set to 20. The plotted using packages in R. Midpoint-root and ladderized were carried out using phytools[72] and ape [73], and ggtree [74] was used for the visualisation. The subgenus information for Betacoronavirus were curated and clades labelled based on consensus of the majority (i.e. if > 85% of the samples in the clade are labelled and have the same subgenus annotation). For labelling the bat-CoVs host genera and species information, a list of host genera and species are curated. Host species with >10 bat-CoV genomes were labelled, followed by host genera with more > 10 bat-CoV genomes. The remaining bats were grouped into a single group “other”.

### 4.4. Gene Relationship Network Graph

Genes identified by PROKKA from each host-set were collated and together with the additional sequences from the BLAST-alignment to the SARS-CoV-2 ref genome as aforementioned, an all-against-all comparison was made with BLAST. This was done with all gene sequences as both the reference and the query as input. A network graph was generated using Graphia Enterprise [75] by treating each gene as a node and generating edges between nodes with significant BLAST alignments. A significant BLAST alignment was defined to have a BLAST score ≥ 60, a query coverage ≥ 80% and a percentage identity ≥ 80%. Components with less than 5 nodes were removed from the graph. The same procedure was carried out using amino acid sequences as input (Figure A1). Where the amino acid sequences were not generated by PROKKA, the matched sequences extracted from BLAST were translated into amino acid sequences, provided that the sequences contained the start and stop codons.

### 4.5. Codon Usage

Codon usage metrics for every gene in the SARS-CoV-2 reference gene catalogue were calculated in all available genome sets. Gene sequence output of the PROKKA and BLAST searches (where correct frame was present) were collated and BLAST searched against the SARS-CoV-2 ref genes; genes that have a BLAST result were included and annotated with the SARS-CoV-2 gene. For each set of genes annotated with an SARS-CoV-2 gene, those substantially shorter than the average (< mean length - 2 standard deviation) were removed from codon usage analysis. Custom Python scripts (available on Github (https://github.com/coronahack2020/final_paper.git) were used to summarise the frequencies of each of the codons. Non-standard codons, start, stop codons were discarded, along with the codon TGG as it is the only codon codinig for tryptophan.

Relative synonymous codon usage was calculated as the ratio of the observed frequency of codon to the expected frequency under the assumption of equal usage between synonymous codons for the same amino acids [76].

### 4.6. Variant Analysis

For this analysis, we aim to highlight naturally occurring and population-wide viable variants, defined as being different to the SARS-CoV-2 ref and have an impact on coding potential. Variant calling was carried out for all available genome sets against the reference SARS-CoV-2 genome released by Ensembl v100 *ASM985889v3.* The allelic counts and variant effect prediction was carried out in order to identify variants with high impact changes (inframe deletion, inframe insertion, frameshift, or stop gain) within or between viruses collected from different host species.

Briefly, multiple genome fasta input files were mapped against the SARS-CoV-2 ref assembly using minimap2 [77] with the following flags (minimap2 -cs -cx asm20 INPUT REF > OUT.paf). The generated PAF (pairwise alignment format) files were subsequently used for variant calling through the paftools.js module in minimap2 (sort -k6,6 -k8,8n OUT.paf I paftools.js call -l 200 -L 200 -q 30 -f REF.fa). Haplotype aware variant consequences were generated using VEP (Variant Effect Predictor) [78] [79]) and BCFtools/csq [80]. The complete set of scripts for this pipeline can be found in https://github.com/coronahack2020/final_paper.git.

### 4.7. Expression Analysis

The RNASeq dataset (n=4) was obtained from the publicly available project PRJCA002326 at National Genomic Data Centre of Beijing Genomics Institute.

The details of the samples can be found in https://bigd.big.ac.cn/bioproject/browse/PRJCA002326. Briefly, total RNA were extracted from broncho-alveolar flush (BALF) samples of two COVID-19 patients treated at the Wuhan University Hospital (Wuhan, China). Ribosomal depeltion was carried out, followed by 150bp pair-end sequencing with an 145bp insert size using Illumina MiSeq. After trimming the raw reads using Trimmomatic v.0.39 [81], a Kallisto index was built based on cDNA fasta obtained from Ensembl v100 *ASM985889v3.* After mapping the read to transcriptome (CDS) level fasta file using Kallisto, the transcript level abundance (TPM) was extracted and visualised in R v.4.0.0 [82].

## Author Contributions

Study conceptualisation, methodology and formal analysis was carried out by Nicholas J Dimonaco (NJD), Barbara B. Shih (BBS) and Mazdak Salavati (MS). Data curation was carried out by NJD. Writing was carried out by NJS, BBS and MS. Visualization was carried out by BBS and MS.

## Acknowledgments

This study was carried out with support from DataBiology, MindStreamAI, University of Edinburgh, The Roslin Institute Royal (Dick) School of Veterinary Studies, Institute of Genetics and Molecular Medicine and University of Aberystwyth. Authors of this manuscript were members of the team who one the 3rd joint position in CORONAHACK2020 virtual hackathon. The full team members were Mazdak Salavati, Barbara B. Shih, Nicholas J. Dimonaco and David A. Parry that contributed equally to the hackathon’s outcome. The prize of the Hackathon sponsored by Slack, Fluidstack, Episode 1, Scan Computers, DataBiology, NVIDIA and MindStreamAI (£500) was used towards publication fees of this manuscript.

NJD was awarded the Rhiannon Powell Science Bursary by the Old Students’ Association of Aberystwyth University in support of his contribution to the manuscript. Please refer to this link for the details of the event: https://www.coronahack.co.uk/ Thanks to Dr Samantha Lycett, Roslin Institute for comments on the manuscript. BBS is supported by a BBSRC Core Capability Grant (BB/CCG1780/1) to the Roslin Institute.

## Abbreviations

The following abbreviations are used in this manuscript:

BLAST: Basic Local Alignment Search Tool
CoV: Coronavirus
DB: DataBiology
E: Envelope
hACE2: human Angiotensin Converting Enzyme
M: Membrane
MERS-CoV: Middle East Respiratory Syndrome Coronavirus N Nucleocapsid
NCBI: National Center for Biotechnology Information
ORF: Open reading frame
PCA: Principle component analysis
PC: Principle component
PROKKA: Rapid Prokaryotic Genome Annotation
RaTG13: SARSr-Ra-BatCoV-RaTG13
RBD: Receptor binding domain
RSCU: Relative synonymous codon usage
SARS: Severe Acute Respiratory Syndrome
S: Spike glycoprotein
ViPR: Virus Pathogen Resource

## Appendix A. Phylogenetic tree on a subset of samples

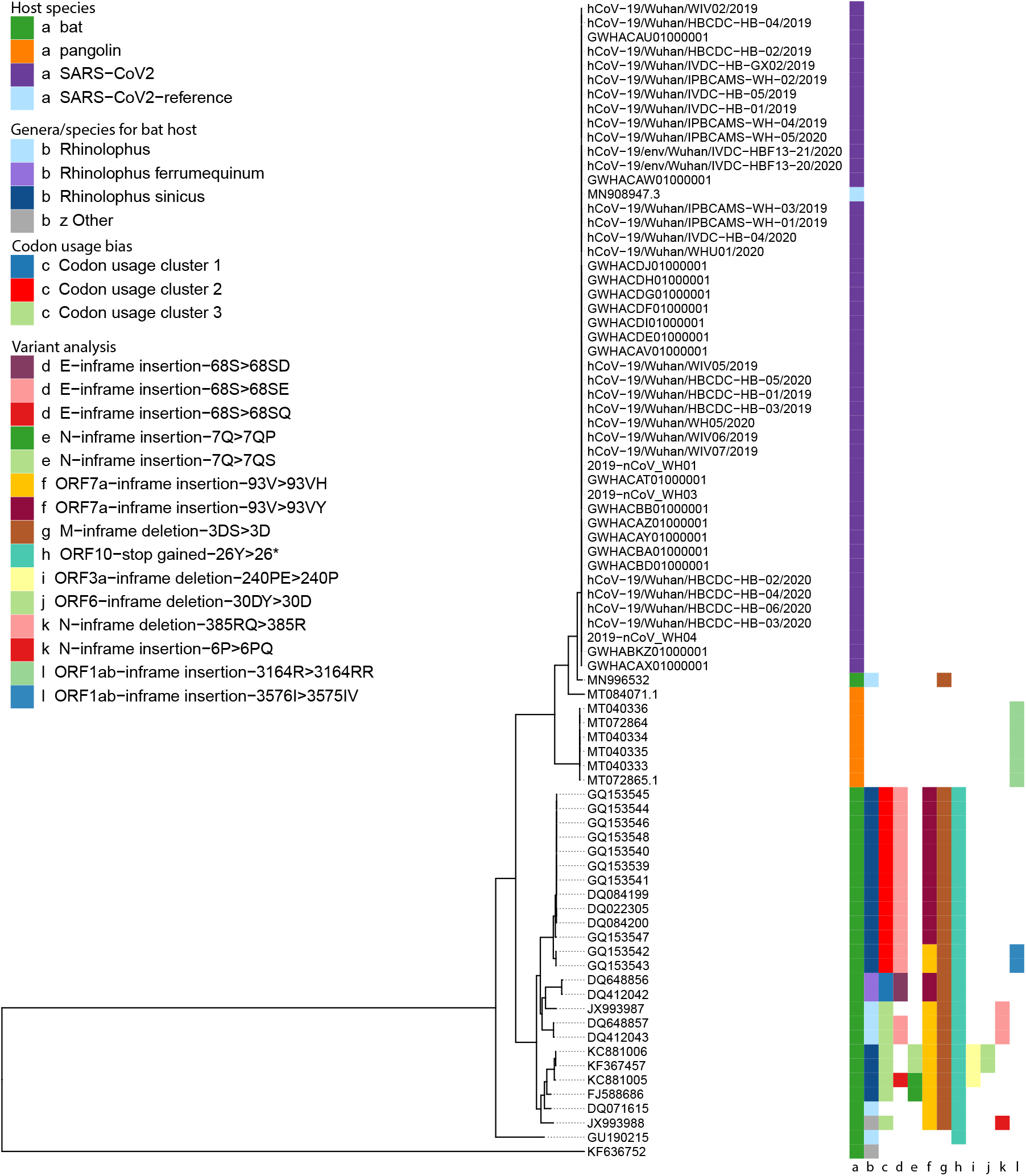

## Appendix B. Genome Annotation Presented by Source

**Table A1.**
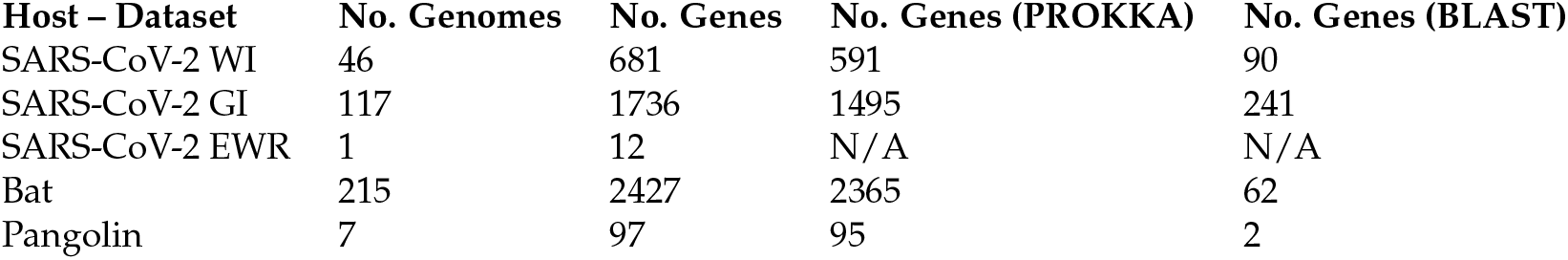
Table containing the total number of genomes and genes for each host-species group. Gene-sets listing number of genes identified by either PROKKA or BLAST. SARS-CoV-2 group names shortened as; WI: Wuhan Isolates, GI: German Isolates, EWR: Ensembl Wuhan Reference. Listed is the total number of all PROKKA genes identified and the number of BLAST genes which matched an Ensembl reference gene with 80% percentage identity.

| Host – Dataset | No. Genomes | No. Genes | No. Genes (PROKKA) | No. Genes (BLAST) |
| --- | --- | --- | --- | --- |
| SARS-CoV-2 WI | 46 | 681 | 591 | 90 |
| SARS-CoV-2 GI | 117 | 1736 | 1495 | 241 |
| SARS-CoV-2 EWR | 1 | 12 | N/A | N/A |
| Bat | 215 | 2427 | 2365 | 62 |
| Pangolin | 7 | 97 | 95 | 2 |

## Appendix C. Gene-gene network graph using amino acid sequences

**Figure A1.**
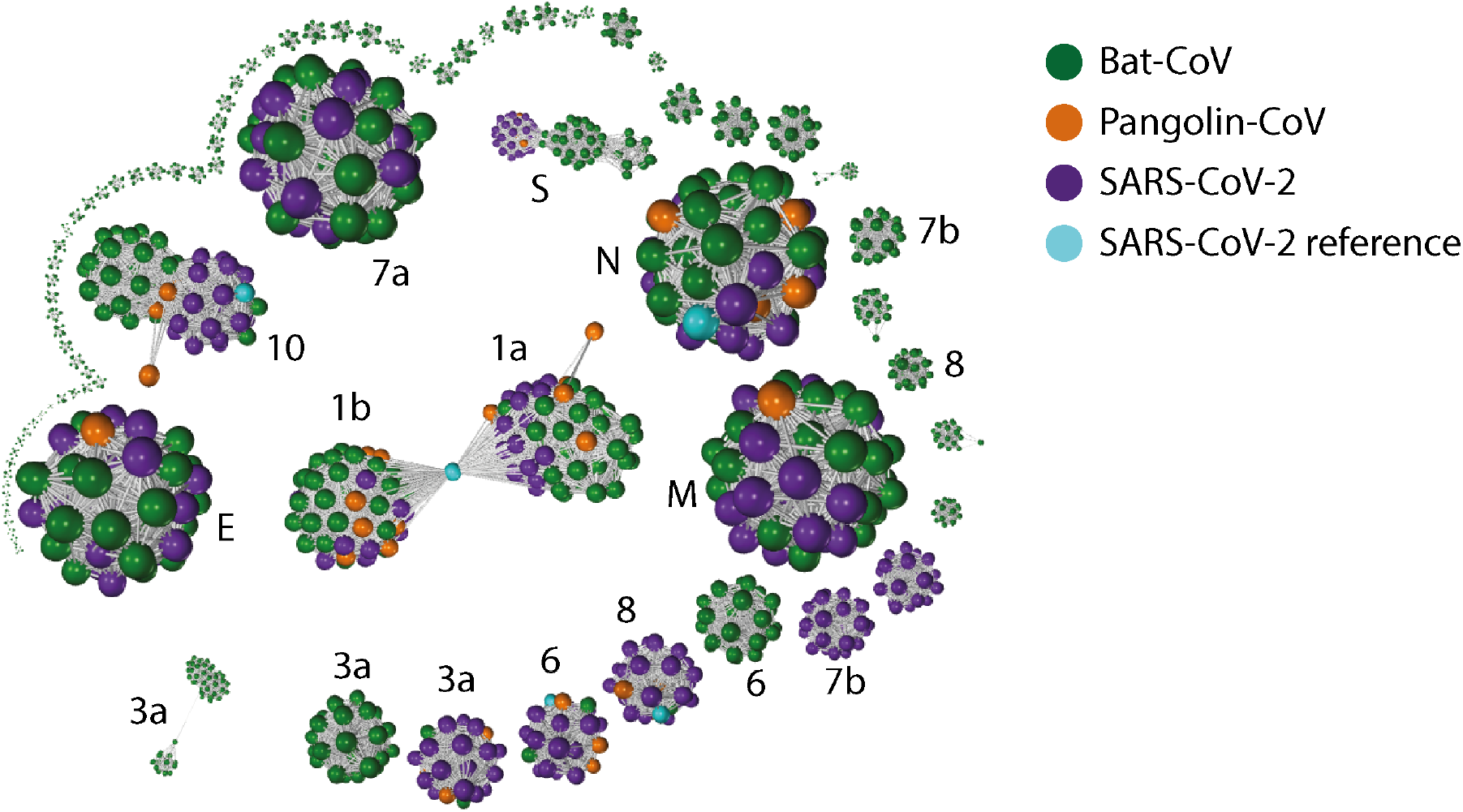
Gene-gene similarity network analysis. Each node represents a amino acid sequence defined by PROKKA or BLAST (ORF10 and E). The nodes were compared against each other using BLAST, and nodes with high similarity (BLAST score ≥ 60 and a query coverage ≥ 80%) were connected with an edge. The network graph is labelled with with SARS-CoV-2 gene names (“ORF” omitted). When the network graph is coloured by host species, genes showing higher degree of variability across species are highlighted. Similar to the network analysis on nucleotide sequences (Figure 2). Genes ORF3a, ORF6, ORF7b, ORF8, ORF10 and S show strong separation between nodes from different species. The degree of separation in ORF1ab are stronger than ORF10 in the nucleic acid network graph; the reverse is true for the amino acid network graph.

## Appendix D. PCA plots based on the RSCU for each gene

**Figure A2.**
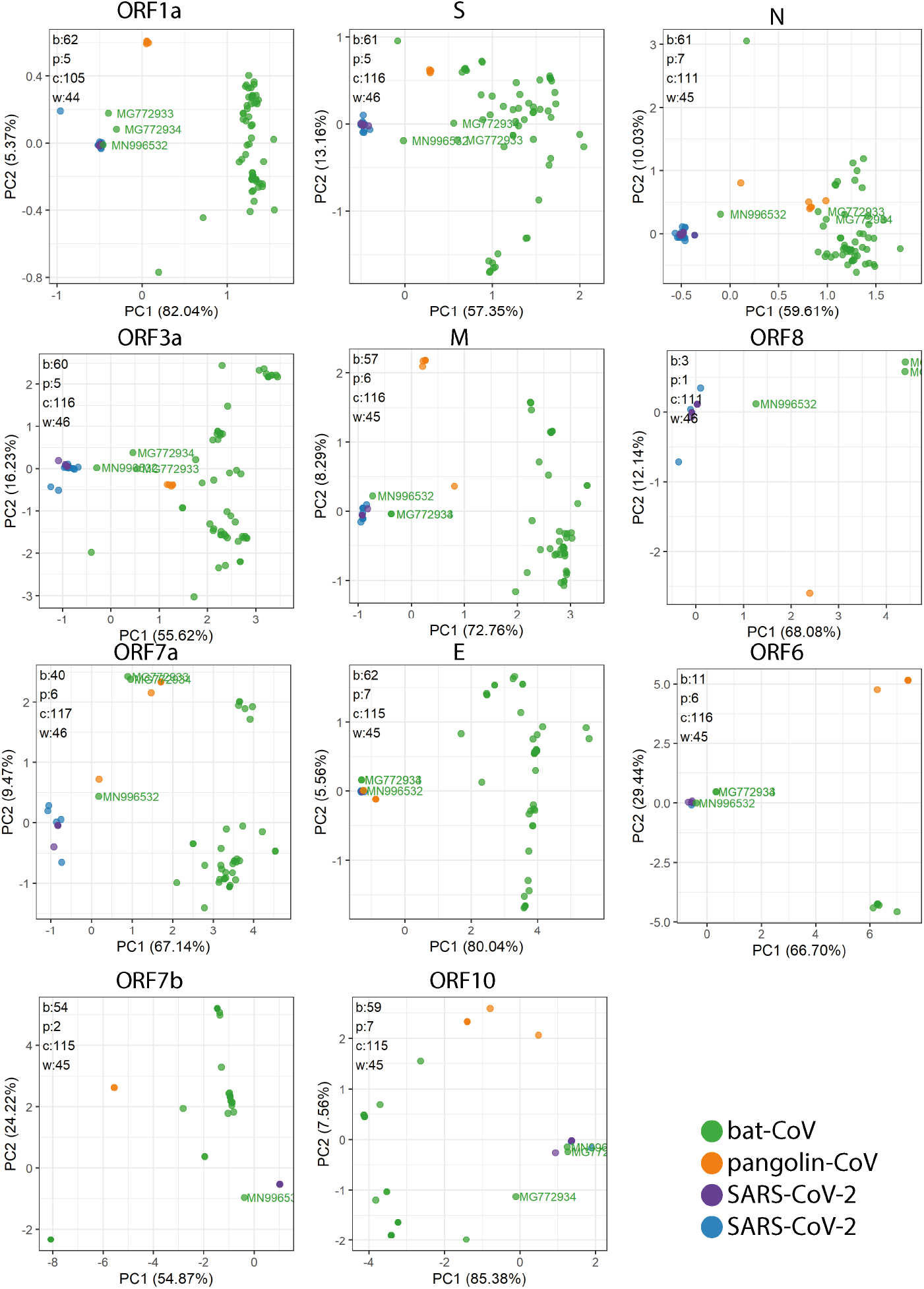
Relative synonymous codon usage (RSCU) was calculated as the ratio of the observed frequency of codon to the expected frequency under the assumption of equal usage between synonymous codons for the same amino acids. This was carried out for each gene. The total number of genomes used in each plot are indicated in the top left corner for bat-CoV (b), pangolin-CoV (p),SARS-CoV-2 Charite dataset (c) and Wuhan dataset (w). As well as a strong separation between bat-CoV and SARS-CoV-2, there is also some separation within bat-CoV for most genes. Whilst we have illustrated the PCA based on RSCU for all genes, the interpretation for some of the shorter genes should be done with caution as they do not encompass the full spectrum of amino acids.

## Appendix E. PCA plots based on the RSCU for each gene

**Figure A3.**
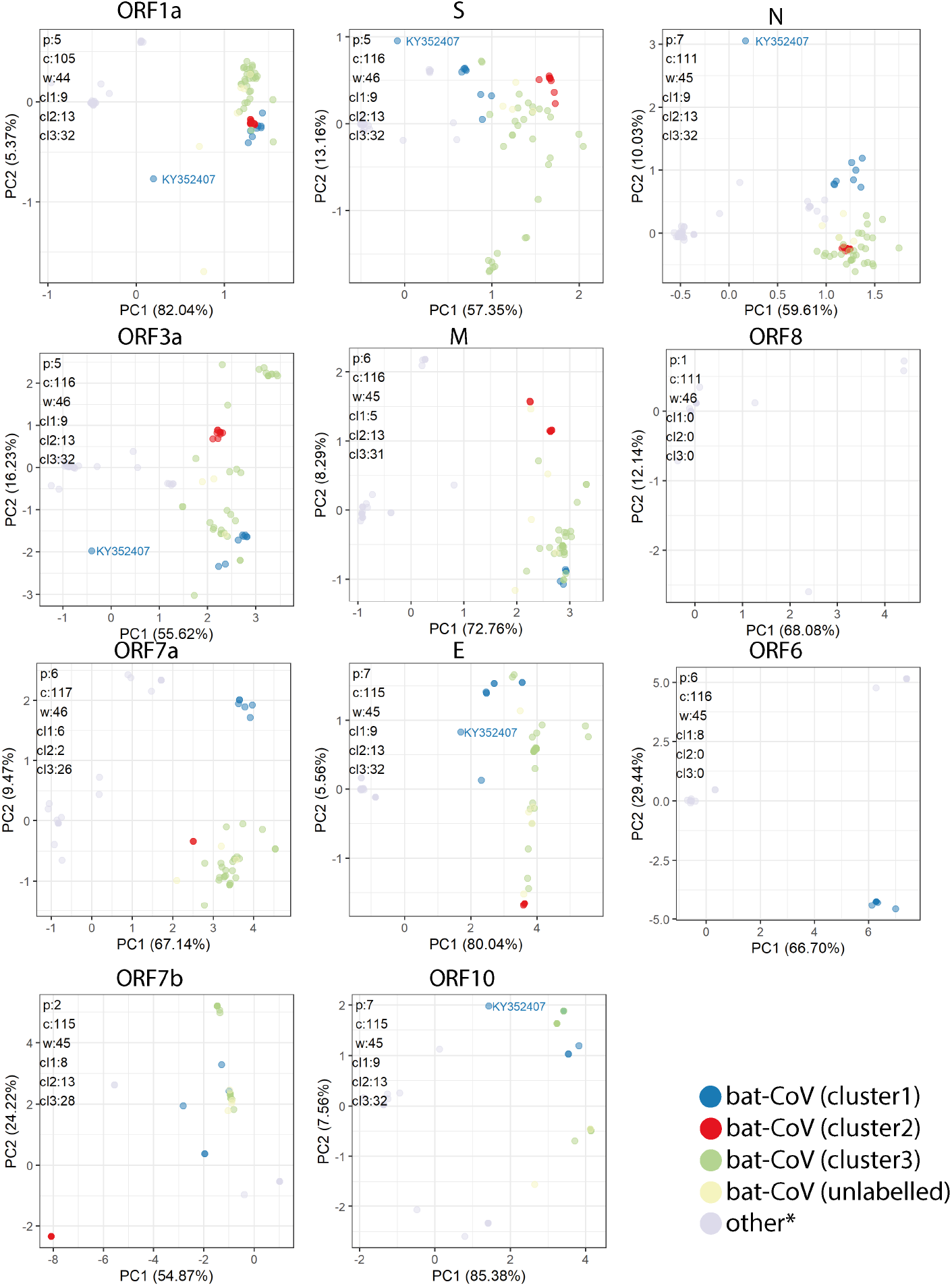
Relative synonymous codon usage (RSCU) was calculated as the ratio of the observed frequency of codon to the expected frequency under the assumption of equal usage between synonymous codons for the same amino acids. This was carried out for each gene. The total number of genomes used in each plot are indicated in the top left corner for pangolin-CoV (p), SARS-CoV-2 Charite dataset (c), Wuhan dataset (w), cluster 1 bat-CoV (cl1), cluster 2 bat-CoV (cl2) and cluster 3 bat-CoV (cl3). The clustering for bat-CoV refers to the k-means clustering performed on PCA of RSCU using multiple genes (Figure 3). As well as a strong separation between bat-CoV and SARS-CoV-2, there is also some separation within bat-CoV for most genes. The clusters seen in RSCU PCA with multiple gene remain together for all genes where the majorities of the genomes are present. Whilst we have illustrated the PCA based on RSCU for all genes, the interpretation for some of the shorter genes should be done with caution as they do not encompass the full spectrum of amino acids.

## Appendix F. RNAseq analysis of SARS-CoV-2-mapping reads

**Figure A4.**
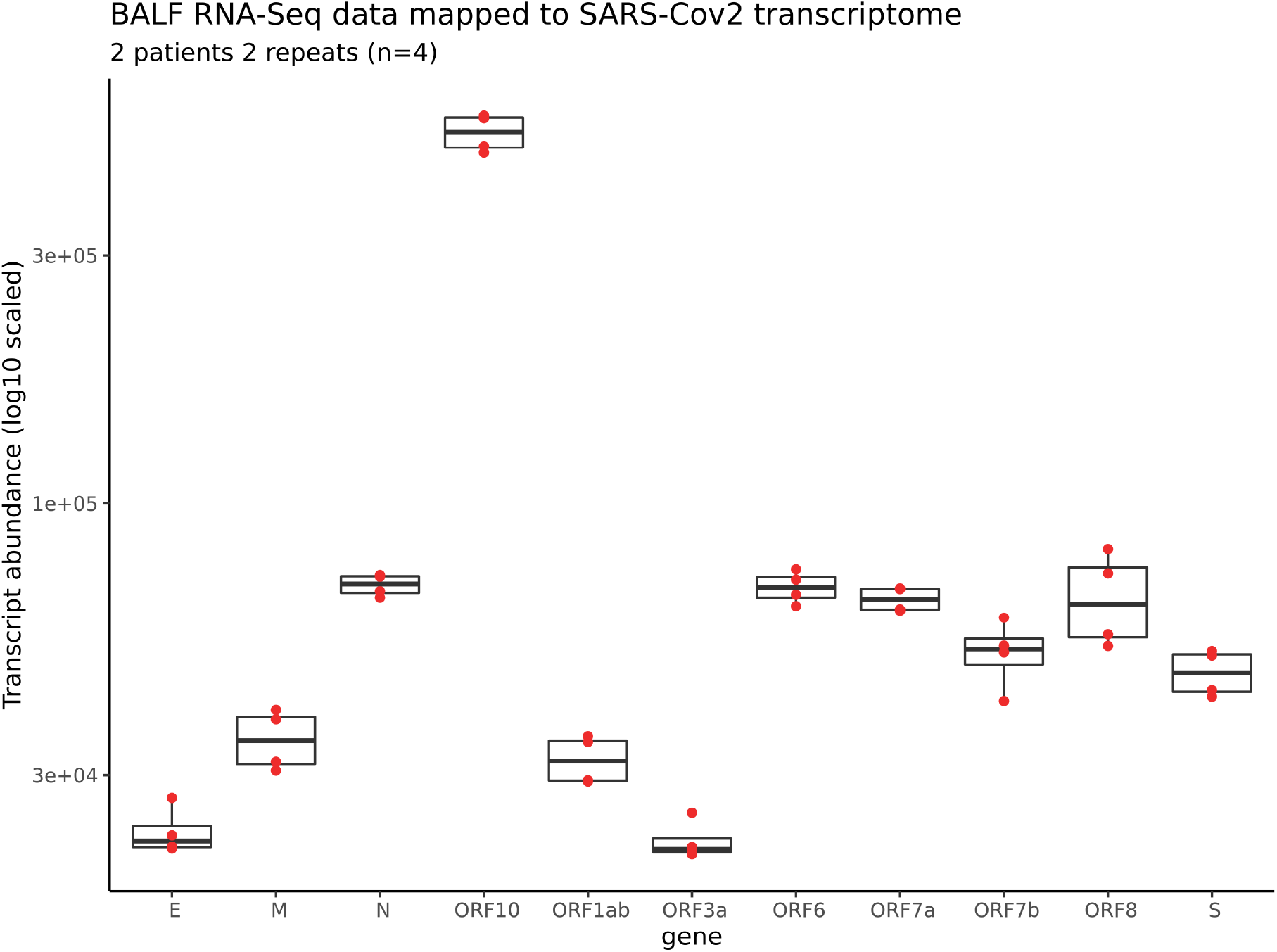
Transcript level expression estimated using Kallisto on SARS-CoV-2 in broncho-alveolar flush (BALF) samples (n=4) collected from 2 patients in Wuhan outbreak. The results is shown here is inaccurate and for record purpose only. This analysis was done during the Hackathon event, during which we had not appreciated the importance of removing reads from the host organism nor did we recognised the lack of distinction between reads mapping to the viral genome or mRNA using this method.

## Appendix G. Synonymous codon ratios

**Figure A5.**
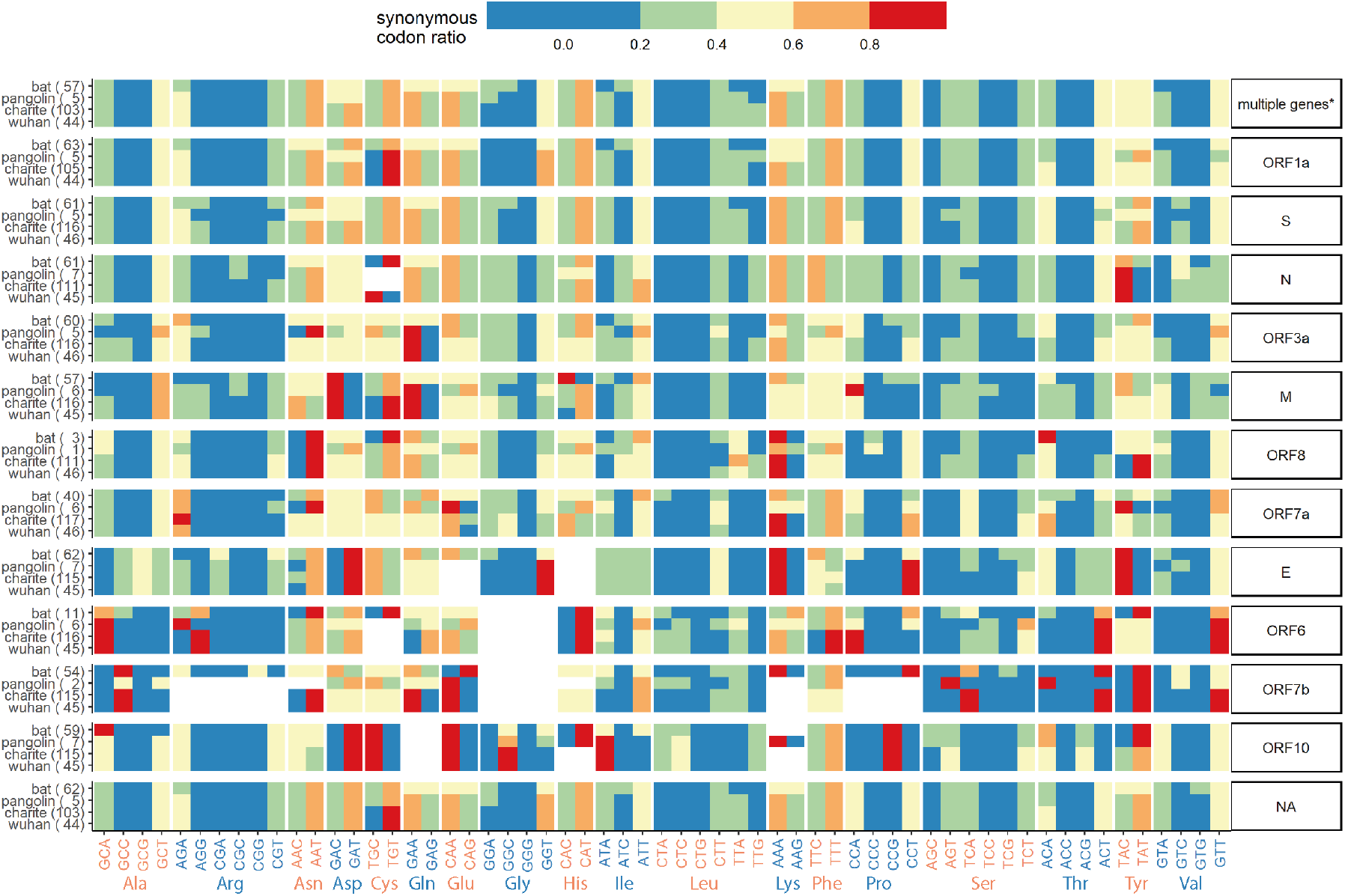
Synonymous codon ratios are the ratio between the number of a given codon divided by the total number of codon coding for the same amino acid. By sorting this ratio in blocks of synonymous codons, this heatmap illustrate the preferential codons for each amino acid for each dataset across all genes. A number of codon usage bias are consistent across most genes and datasets. For instance, GCT is preferentially used for Alanine and GTT for Valine. On the whole, there seem to be less of a preferential codon use for bat, especially in longer genes or when multiple genes are accounted for, as per indicated by the higher frequency of more evenly distributed codon within each amino acid (i.e. for the bat dataset, the heatmap colours are of a similar level within each amino acid). Codons with GCs are generally underrepresented, such as in Arg (Arginine), Pro (Proline) and Ser (Serine). * The values in this row is generated by combing codons from multiple genes, E, N, S, ORF1ab, ORF3a, ORF10.

## Appendix H. Combined variant analysis

**Figure A6.**
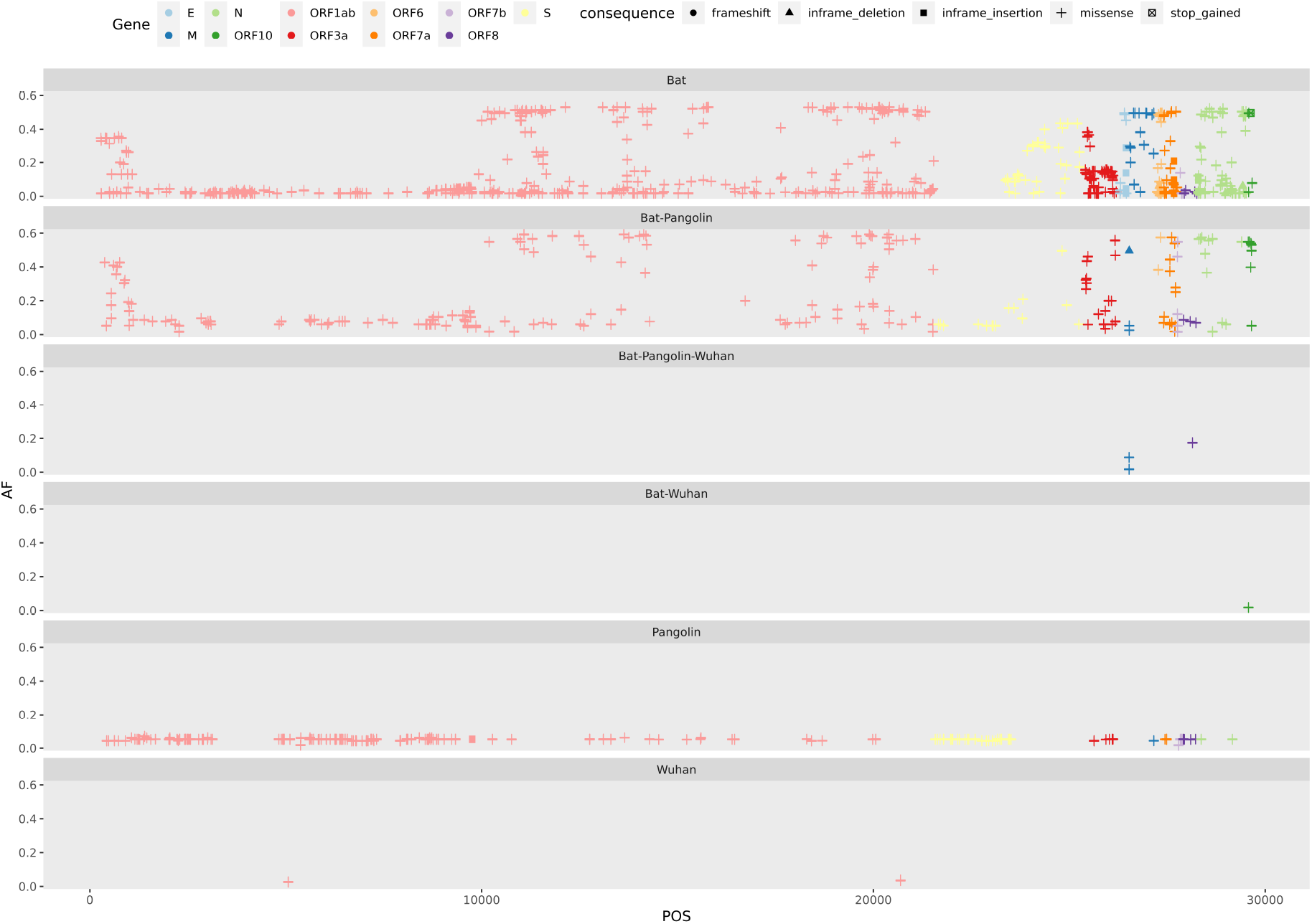
The coordinate map of all variants called against the human reference SARS-Cov-2 genome. Each horizontal track shows the variants present in the host-specie group. The colours shows the gene annotation origin of the variant and the shape consequence

© 2020 by the authors. Submitted to *Viruses* for possible open access publication under the terms and conditions of the Creative Commons Attribution (CC BY) license (http://creativecommons.org/licenses/by/4.0/).

## References

1. Patz, J.A.; Graczyk, T.K.; Geller, N.; Vittor, A.Y. Effects of environmental change on emerging parasitic diseases. International journal for parasitology 2000, 30, 1395–1405.

2. Madhav, N.; Oppenheim, B.; Gallivan, M.; Mulembakani, P.; Rubin, E.; Wolfe, N. Pandemics: risks, impacts, and mitigation; The International Bank for Reconstruction and Development / The World Bank, 2017.

3. of the International, C.S.G.; others. The species Severe acute respiratory syndrome-related coronavirus: classifying 2019-nCoV and naming it SARS-CoV-2. Nature Microbiology 2020, 5, 536.

4. Weiss, S.R. Forty years with coronaviruses. Journal of Experimental Medicine 2020, 217.

5. Perlman, S.; Netland, J. Coronaviruses post-SARS: update on replication and pathogenesis. Nature reviews microbiology 2009, 7, 439–450.

6. Amer, H.; Alqahtani, A.S.; Alzoman, H.; Aljerian, N.; Memish, Z.A. Unusual presentation of Middle East respiratory syndrome coronavirus leading to a large outbreak in Riyadh during 2017. American journal of infection control 2018, 46, 1022–1025.

7. Hung, L.S. The SARS epidemic in Hong Kong: what lessons have we learned? Journal of the Royal Society of Medicine 2003, 96, 374–378.

8. Zhu, Z.; Lian, X.; Su, X.; Wu, W.; Marraro, G.A.; Zeng, Y. From SARS and MERS to COVID-19: a brief summary and comparison of severe acute respiratory infections caused by three highly pathogenic human coronaviruses. Respiratory research 2020, 21, 1–14.

9. Boni, M.F.; Lemey, P.; Jiang, X.; Lam, T.T.Y.; Perry, B.; Castoe, T.; Rambaut, A.; Robertson, D.L. Evolutionary origins of the SARS-CoV-2 sarbecovirus lineage responsible for the COVID-19 pandemic. bioRxiv 2020, [https://www.biorxiv.org/content/early/2020/03/31/2020.03.30.015008.full.pdf]. doi:10.1101/2020.03.30.015008.

10. Zhang, Y.Z.; Holmes, E.C. A Genomic Perspective on the Origin and Emergence of SARS-CoV-2. Cell 2020, 181, 223 – 227. doi:https://doi.org/10.1016/j.cell.2020.03.035.

11. Jitobaom, K.; Phakaratsakul, S.; Sirihongthong, T.; Chotewutmontri, S.; Suriyaphol, P.; Suptawiwat, O.; Auewarakul, P. Codon usage similarity between viral and some host genes suggests a codon-specific translational regulation. Heliyon 2020, 6, e03915.

12. Kumar, N.; Kulkarni, D.D.; Lee, B.; Kaushik, R.; Bhatia, S.; Sood, R.; Pateriya, A.K.; Bhat, S.; Singh, V.P. Evolution of codon usage bias in Henipaviruses is governed by natural selection and is host-specific. Viruses 2018, 10, 604.

13. Chen, F.; Wu, P.; Deng, S.; Zhang, H.; Hou, Y.; Hu, Z.; Zhang, J.; Chen, X.; Yang, J.R. Dissimilation of synonymous codon usage bias in virus–host coevolution due to translational selection. Nature ecology & evolution 2020, pp. 1–12.

14. Elbe, S.; Buckland-Merrett, G.; falkename, t.; thistoo, a. Data, disease and diplomacy: GISAID’s innovative contribution to global health. Global Challenges 2017, 1, 33–46.

15. Yates, A.D.; Achuthan, P.; Akanni, W.; Allen, J.; Allen, J.; Alvarez-Jarreta, J.; Amode, M.R.; Armean, I.M.; Azov, A.G.; Bennett, R.; others. Ensembl 2020. Nucleic acids research 2020, 48, D682–D688.

16. Seemann, T. Prokka: rapid prokaryotic genome annotation. Bioinformatics 2014, 30, 2068–2069.

17. Altschul, S.F.; Gish, W.; Miller, W.; Myers, E.W.; Lipman, D.J. Basic local alignment search tool. Journal of molecular biology 1990, 215, 403–410.

18. Baranov, P.V.; Henderson, C.M.; Anderson, C.B.; Gesteland, R.F.; Atkins, J.F.; Howard, M.T. Programmed ribosomal frameshifting in decoding the SARS-CoV genome. Virology 2005, 332, 498–510.

19. Robertson, M.P.; Igel, H.; Baertsch, R.; Haussler, D.; Ares Jr, M.; Scott, W.G. The structure of a rigorously conserved RNA element within the SARS virus genome. PLoS Biol 2004, 3, e5.

20. Tengs, T.; Jonassen, C.M. Distribution and evolutionary history of the mobile genetic element s2m in coronaviruses. Diseases 2016, 4, 27.

21. Tengs, T.; Delwiche, C.F.; Jonassen, C.M. A mobile genetic element in the SARS-CoV-2 genome is shared with multiple insect species. bioRxiv 2020.

22. Lopes, L.R.; de Mattos Cardillo, G.; Paiva, P.B. Molecular evolution and phylogenetic analysis of SARS-CoV-2 and hosts ACE2 protein suggest Malayan pangolin as intermediary host. Brazilian Journal of Microbiology 2020, pp. 1–7.

23. Fahmi, M.; Kubota, Y.; Ito, M. Nonstructural proteins NS7b and NS8 are likely to be phylogenetically associated with evolution of 2019-nCoV. Infection, Genetics and Evolution 2020, 81, 104272.

24. Xiao, K.; Zhai, J.; Feng, Y.; Zhou, N.; Zhang, X.; Zou, J.J.; Li, N.; Guo, Y.; Li, X.; Shen, X.; others. Isolation of SARS-CoV-2-related coronavirus from Malayan pangolins. Nature 2020, pp. 1–4.

25. Li, Y.; Yang, X.; Wang, N.; Wang, H.; Yin, B.; Yang, X.; Jiang, W. The divergence between SARS-CoV-2 and RaTG13 might be overestimated due to the extensive RNA modification. Future Virology 2020.

26. Lau, S.K.; Luk, H.K.; Wong, A.C.; Li, K.S.; Zhu, L.; He, Z.; Fung, J.; Chan, T.T.; Fung, K.S.; Woo, P.C. Possible bat origin of severe acute respiratory syndrome coronavirus 2. Emerging infectious diseases 2020, 26, 1542.

27. Malaiyan, J.; Arumugam, S.; Mohan, K.; Radhakrishnan, G.G. An update on origin of SARS-CoV-2: Despite closest identity, bat (RaTG13) and Pangolin derived Coronaviruses varied in the critical binding site and O-linked glycan residues. Journal of medical virology 2020.

28. Li, W.; Shi, Z.; Yu, M.; Ren, W.; Smith, C.; Epstein, J.H.; Wang, H.; Crameri, G.; Hu, Z.; Zhang, H.; others. Bats are natural reservoirs of SARS-like coronaviruses. Science 2005, 310, 676–679.

29. Banerjee, A.; Kulcsar, K.; Misra, V.; Frieman, M.; Mossman, K. Bats and coronaviruses. Viruses 2019, 11, 41.

30. Su, S.; Wong, G.; Shi, W.; Liu, J.; Lai, A.C.; Zhou, J.; Liu, W.; Bi, Y.; Gao, G.F. Epidemiology, genetic recombination, and pathogenesis of coronaviruses. Trends in microbiology 2016, 24, 490–502.

31. Yi, H. 2019 novel coronavirus is undergoing active recombination. Clinical Infectious Diseases 2020.

32. Koyama, T.; Platt, D.; Parida, L. Variant analysis of SARS-CoV-2 genomes. Bulletin of the World Health Organization 2020, 98.

33. Zhang, T.; Wu, Q.; Zhang, Z. Probable pangolin origin of SARS-CoV-2 associated with the COVID-19 outbreak. Current Biology 2020.

34. Ceraolo, C.; Giorgi, F.M. Genomic variance of the 2019-nCoV coronavirus. Journal of medical virology 2020, 92, 522–528.

35. Pereira, F. Evolutionary dynamics of the SARS-CoV-2 ORF8 accessory gene. Infection, Genetics and Evolution 2020, 85, 104525.

36. Mohammad, S.; Bouchama, A.; Mohammad Alharbi, B.; Rashid, M.; Saleem Khatlani, T.; Gaber, N.S.; Malik, S.S. SARS-CoV-2 ORF8 and SARS-CoV ORF8ab: Genomic Divergence and Functional Convergence. Pathogens 2020, 9, 677.

37. Li, J.Y.; Liao, C.H.; Wang, Q.; Tan, Y.J.; Luo, R.; Qiu, Y.; Ge, X.Y. The ORF6, ORF8 and nucleocapsid proteins of SARS-CoV-2 inhibit type I interferon signaling pathway. Virus research 2020, 286, 198074.

38. Wong, H.H.; Fung, T.S.; Fang, S.; Huang, M.; Le, M.T.; Liu, D.X. Accessory proteins 8b and 8ab of severe acute respiratory syndrome coronavirus suppress the interferon signaling pathway by mediating ubiquitin-dependent rapid degradation of interferon regulatory factor 3. Virology 2018, 515, 165–175.

39. Hachim, A.; Kavian, N.; Cohen, C.A.; Chin, A.W.; Chu, D.K.; Mok, C.K.; Tsang, O.T.; Yeung, Y.C.; Perera, R.A.; Poon, L.L.; others. ORF8 and ORF3b antibodies are accurate serological markers of early and late SARS-CoV-2 infection. Technical report, Nature Publishing Group, 2020.

40. Lau, S.K.; Feng, Y.; Chen, H.; Luk, H.K.; Yang, W.H.; Li, K.S.; Zhang, Y.Z.; Huang, Y.; Song, Z.Z.; Chow, W.N.; others. Severe acute respiratory syndrome (SARS) coronavirus ORF8 protein is acquired from SARS-related coronavirus from greater horseshoe bats through recombination. Journal of virology 2015, 89, 10532–10547.

41. Consortium, C.S.M.E.; others. Molecular evolution of the SARS coronavirus during the course of the SARS epidemic in China. Science 2004, 303, 1666–1669.

42. Wrapp, D.; Wang, N.; Corbett, K.S.; Goldsmith, J.A.; Hsieh, C.L.; Abiona, O.; Graham, B.S.; McLellan, J.S. Cryo-EM structure of the 2019-nCoV spike in the prefusion conformation. Science 2020, 367, 1260–1263, [https://science.sciencemag.org/content/367/6483/1260.full.pdf]. doi:10.1126/science.abb2507.

43. Wu, K.; Chen, L.; Peng, G.; Zhou, W.; Pennell, C.A.; Mansky, L.M.; Geraghty, R.J.; Li, F. A virus-binding hot spot on human angiotensin-converting enzyme 2 is critical for binding of two different coronaviruses. Journal of virology 2011, 85, 5331–5337.

44. Hoffmann, M.; Kleine-Weber, H.; Schroeder, S.; Krüger, N.; Herrler, T.; Erichsen, S.; Schiergens, T.S.; Herrler, G.; Wu, N.H.; Nitsche, A.; others. SARS-CoV-2 cell entry depends on ACE2 and TMPRSS2 and is blocked by a clinically proven protease inhibitor. Cell 2020.

45. Baranowski, E.; Ruiz-Jarabo, C.M.; Domingo, E. Evolution of cell recognition by viruses. Science 2001, 292, 1102–1105.

46. Baranowski, E.; Ruiz-Jarabo, C.M.; Pariente, N.; Verdaguer, N.; Domingo, E. Evolution of cell recognition by viruses: a source of biological novelty with medical implications. Advances in virus research 2003, 62, 19.

47. Qu, X.X.; Hao, P.; Song, X.J.; Jiang, S.M.; Liu, Y.X.; Wang, P.G.; Rao, X.; Song, H.D.; Wang, S.Y.; Zuo, Y.; others. Identification of two critical amino acid residues of the severe acute respiratory syndrome coronavirus spike protein for its variation in zoonotic tropism transition via a double substitution strategy. Journal of Biological Chemistry 2005, 280, 29588–29595.

48. Rice, A.M.; Morales, A.C.; Ho, A.T.; Mordstein, C.; Mühlhausen, S.; Watson, S.; Cano, L.; Young, B.; Kudla, G.; Hurst, L.D. Evidence for strong mutation bias towards, and selection against, U content in SARS-CoV-2: implications for vaccine design. Molecular biology and evolution 2020, p. msaa188. PMC7454790[pmcid], doi:10.1093/molbev/msaa188.

49. Gu, H.; Chu, D.K.; Peiris, M.; Poon, L.L. Multivariate analyses of codon usage of SARS-CoV-2 and other betacoronaviruses. Virus Evolution 2020, 6, veaa032.

50. Nambou, K.; Anakpa, M. Deciphering the co-adaptation of codon usage between respiratory coronaviruses and their human host uncovers candidate therapeutics for COVID-19. Infection, Genetics and Evolution 2020, 85, 104471.

51. Alonso, A.M.; Diambra, L. SARS-CoV-2 Codon Usage Bias Downregulates Host Expressed Genes With Similar Codon Usage. Frontiers in Cell and Developmental Biology 2020, 8, 831. doi:10.3389/fcell.2020.00831.

52. Digard, P.; Lee, H.M.; Sharp, C.; Grey, F.; Gaunt, E.R. Intra-genome variability in the dinucleotide composition of SARS-CoV-2. bioRxiv 2020.

53. Sanjuán, R.; Domingo-Calap, P. Mechanisms of viral mutation. Cellular and molecular life sciences 2016, 73, 4433–4448.

54. Smith, E.C.; Sexton, N.R.; Denison, M.R. Thinking outside the triangle: replication fidelity of the largest RNA viruses. Annual Review of Virology 2014, 1, 111–132.

55. Hassan, S.S.; Attrish, D.; Ghosh, S.; Choudhury, P.P.; Uversky, V.N.; Uhal, B.D.; Lundstrom, K.; Rezaei, N.; Aljabali, A.A.; Seyran, M.; others. Notable sequence homology of the ORF10 protein introspects the architecture of SARS-COV-2. bioRxiv 2020.

56. Kim, D.; Lee, J.Y.; Yang, J.S.; Kim, J.W.; Kim, V.N.; Chang, H. The architecture of SARS-CoV-2 transcriptome. Cell 2020.

57. Liu, T.; Jia, P.; Fang, B.; Zhao, Z. Differential expression of viral transcripts from single-cell RNA sequencing of moderate and severe COVID-19 patients and its implications for case severity. Frontiers in Microbiology 2020, 11, 2568.

58. Subissi, L.; Posthuma, C.C.; Collet, A.; Zevenhoven-Dobbe, J.C.; Gorbalenya, A.E.; Decroly, E.; Snijder, E.J.; Canard, B.; Imbert, I. One severe acute respiratory syndrome coronavirus protein complex integrates processive RNA polymerase and exonuclease activities. Proceedings of the National Academy of Sciences 2014, 111, E3900–E3909.

59. Schoeman, D.; Fielding, B.C. Coronavirus envelope protein: current knowledge. Virology journal 2019, 16, 1–22.

60. DeDiego, M.L.; Álvarez, E.; Almazán, F.; Rejas, M.T.; Lamirande, E.; Roberts, A.; Shieh, W.J.; Zaki, S.R.; Subbarao, K.; Enjuanes, L. A severe acute respiratory syndrome coronavirus that lacks the E gene is attenuated in vitro and in vivo. Journal of virology 2007, 81, 1701–1713.

61. Teoh, K.T.; Siu, Y.L.; Chan, W.L.; Schlüter, M.A.; Liu, C.J.; Peiris, J.M.; Bruzzone, R.; Margolis, B.; Nal, B. The SARS coronavirus E protein interacts with PALS1 and alters tight junction formation and epithelial morphogenesis. Molecular biology of the cell 2010, 21, 3838–3852.

62. Hassan, S.S.; Choudhury, P.P.; Roy, B. SARS-CoV2 envelope protein: non-synonymous mutations and its consequences. Genomics 2020.

63. Taylor, J.K.; Coleman, C.M.; Postel, S.; Sisk, J.M.; Bernbaum, J.G.; Venkataraman, T.; Sundberg, E.J.; Frieman, M.B. Severe acute respiratory syndrome coronavirus ORF7a inhibits bone marrow stromal antigen 2 virion tethering through a novel mechanism of glycosylation interference. Journal of virology 2015, 89, 11820–11833.

64. Cong, Y.; Ulasli, M.; Schepers, H.; Mauthe, M.; V’kovski, P.; Kriegenburg, F.; Thiel, V.; de Haan, C.A.; Reggiori, F. Nucleocapsid protein recruitment to replication-transcription complexes plays a crucial role in coronaviral life cycle. Journal of virology 2020, 94.

65. Weber, S.; Ramirez, C.; Doerfler, W. Signal hotspot mutations in SARS-CoV-2 genomes evolve as the virus spreads and actively replicates in different parts of the world. Virus Research 2020, 289, 198170.

66. Pickett, B.E.; Sadat, E.L.; Zhang, Y.; Noronha, J.M.; Squires, R.B.; Hunt, V.; Liu, M.; Kumar, S.; Zaremba, S.; Gu, Z.; others. ViPR: an open bioinformatics database and analysis resource for virology research. Nucleic acids research 2012, 40, D593–D598.

67. Coordinators, N.R. Database resources of the national center for biotechnology information. Nucleic acids research 2018, 46, D8.

68. Dinman, J.D. Programmed–1 Ribosomal Frameshifting in SARS Coronavirus. In Molecular Biology of the SARS-Coronavirus; Springer, 2010; pp. 63–72.

69. Hyatt, D.; Chen, G.L.; LoCascio, P.F.; Land, M.L.; Larimer, F.W.; Hauser, L.J. Prodigal: prokaryotic gene recognition and translation initiation site identification. BMC bioinformatics 2010, 11, 119.

70. Sievers, F.; Higgins, D.G. Clustal Omega for making accurate alignments of many protein sequences. Protein Science 2018, 27, 135–145.

71. Stamatakis, A. RAxML version 8: a tool for phylogenetic analysis and post-analysis of large phylogenies. Bioinformatics 2014, 30, 1312–1313.

72. Revell, L.J. phytools: An R package for phylogenetic comparative biology (and other things). Methods in Ecology and Evolution 2012, 3, 217–223.

73. Paradis, E.; Schliep, K. ape 5.0: an environment for modern phylogenetics and evolutionary analyses in R. Bioinformatics 2019, 35, 526–528.

74. Yu, G. Using ggtree to Visualize Data on Tree-Like Structures. Current protocols in bioinformatics 2020, 69, e96.

75. Freeman, T.; Horsewell, S.; Patir, A.; Harling-Lee, J.; Regan, T.; Shih, B.B.; Prendergast, J.; Hume, D.A.; Angus, T. Graphia: A platform for the graph-based visualisation and analysis of complex data. bioRxiv 2020. doi:10.1101/2020.09.02.279349.

76. Sharp, P.M.; Tuohy, T.M.; Mosurski, K.R. Codon usage in yeast: cluster analysis clearly differentiates highly and lowly expressed genes. Nucleic acids research 1986, 14, 5125–5143.

77. Li, H. Minimap2: pairwise alignment for nucleotide sequences. Bioinformatics 2018, 34, 3094–3100, [https://academic.oup.com/bioinformatics/article-pdf/34/18/3094/25731859/bty191.pdf]. doi:10.1093/bioinformatics/bty191.

78. McLaren, W.; Gil, L.; Hunt, S.E.; Riat, H.S.; Ritchie, G.R.S.; Thormann, A.; Flicek, P.; Cunningham, F. The Ensembl Variant Effect Predictor. Genome Biology 2016, 17, 122. doi:10.1186/s13059-016-0974-4.

79. den Dunnen, J.T.; Dalgleish, R.; Maglott, D.R.; Hart, R.K.; Greenblatt, M.S.; McGowan-Jordan, J.; Roux, A.F.; Smith, T.; Antonarakis, S.E.; Taschner, P.E. HGVS Recommendations for the Description of Sequence Variants: 2016 Update. Human Mutation 2016, 37, 564–569, [https://onlinelibrary.wiley.com/doi/pdf/10.1002/humu.22981]. doi:10.1002/humu.22981.

80. Danecek, P.; McCarthy, S.A. BCFtools/csq: haplotype-aware variant consequences. Bioinformatics 2017, 33, 2037–2039, [https://academic.oup.com/bioinformatics/article-pdf/33/13/2037/25155875/btx100.pdf]. doi:10.1093/bioinformatics/btx100.

81. Bolger, A.M.; Lohse, M.; Usadel, B. Trimmomatic: a flexible trimmer for Illumina sequence data. Bioinformatics (Oxford, England) 2014, 30, 2114–20. doi:10.1093/bioinformatics/btu170.

82. R Core Team. R: A language and environment for statistical computing. R Foundation for Statistical Computing 2020.

